# Regulation of Steady State Ribosomal Transcription in *Mycobacterium tuberculosis*: Intersection of Sigma Subunits, Superhelicity, and Transcription Factors

**DOI:** 10.1101/2025.02.24.639987

**Authors:** Ana Ruiz Manzano, Drake Jensen, Eric A. Galburt

## Abstract

The regulation of ribosomal RNA (rRNA) is closely tied to nutrient availability, growth phase, and global gene expression, serving as a key factor in bacterial adaptability and pathogenicity. *Mycobacterium tuberculosis (Mtb*) stands out from other species with a single ribosomal operon controlled by two promoters: *rrnA*P3 and *rrnA*P1 and a high ratio of sigma (σ) factors to genome size. While the primary σ factor σ^A^ is known to drive ribosomal transcription, the alternative σ factor σ^B^ has been proposed to contribute to the transcription of housekeeping genes, including rRNA under a range of conditions. However, σ^B^’s precise role remains unclear. Here, we quantify steady-state rates in reconstituted transcription reactions and establish that σ^A^-mediated transcription from *rrnA*P3 dominates rRNA production by almost two orders of magnitude with minimal contributions from σ^B^ holoenzymes and/or *rrnA*P1 under all conditions tested. We measure and compare the kinetics of individual initiation steps for both holoenzymes which, taken together with the steady-state rate measurements, lead us to a model where σ^B^ holoenzymes exhibit slower DNA unwinding and slower holoenzyme recycling. Our data further demonstrate that the transcription factors CarD and RbpA reverse or buffer the stimulatory effect of negative superhelicity on σ^A^ and σ^B^ holoenzymes respectively. Lastly, we show that a major determinant of σ^A^’s increased activity is due to its N-terminal 205 amino acids. Taken together, our data reveal the intricate interplay of promoter sequence, σ factor identity, DNA superhelicity, and transcription factors in shaping transcription initiation kinetics and, by extension, the steady-state rates of rRNA production in *Mtb*.

## INTRODUCTION

Roughly one-fourth of the world population has been infected with *Mycobacterium tuberculosis* (*Mtb*), the causative agent of Tuberculosis (TB) disease, which remains one of the leading causes of death worldwide (1). While TB rates in the United States are relatively low, an estimated 13 million individuals live with latent TB infections (2), characterized by an asymptomatic, immune-controlled state (3). Without treatment, approximately 10% of those with latent infection will experience reactivation, a hallmark that distinguishes *Mtb* from many other infectious organisms (1). *Mtb*’s success lies in its ability to sense and adapt to changing environments during infection, a process mainly controlled at the transcriptional level. This adaptation relies on multiple regulatory mechanisms, including the activity of sigma (σ) factors, global transcription regulators like CarD and RbpA, and changes in DNA topology.

Gene expression in bacteria is primarily regulated at transcription initiation, beginning with the formation of the RNA polymerase (RNAP) holoenzyme, which requires the binding of the core RNAP to a σ factor. This complex recognizes promoter DNA, initiating gene expression programs that enable the bacterium to respond to different conditions. *Mtb* has 13 σ factors (4), giving it the highest ratio of σ factors to genome size among human pathogens (5). These σ factors orchestrate gene expression during initial infection, adaptation to stress, and the transition from latent to active disease (6). Among them, σ^A^ is the essential housekeeping σ factor (7–9) in *Mtb* and is analogous to *Escherichia coli* σ^70^ (10, 11). *Mtb* σ^A^ holoenzyme is responsible for transcribing genes required for exponential growth, including those for RNA polymerase, ribosomes, and metabolism (9, 12). In contrast, *Mtb* σ^B^, a non-essential σ factor, plays a key role in stress responses and survival, yet also exhibits activity during exponential growth (13). *Mtb* σ^A^ and σ^B^ share highly homologous C-terminal regions (∼63% identity), including those subregions involved in recognizing the -10 and -35 promoter elements (4, 14–16). Unlike *E. coli*, where σ^S^ (the equivalent of σ^B^) is tightly regulated (17–21), σ^A^ and σ^B^ holoenzymes coexist during exponential phase in *Mtb* (7, 22–24). RNA-seq (9) and ChIP-seq (25) data suggest that the σ^B^ holoenzyme regulates hundreds of genes, including rRNA and housekeeping genes, traditionally thought to be under the exclusive control of σ^A^ (23).

Beyond σ factors, transcription in *Mtb* is further modulated by global regulators like CarD and RbpA. *Mtb* CarD and RbpA are essential global transcription factors required under nutrient-rich conditions and upregulated in response to stress (26–30). Both factors directly interact with RNAP rather than binding to specific DNA sequences (27, 31, 32), and as such, they are often considered ubiquitous members of the initiation complex (33–35). CarD was originally identified as a regulator of rRNA synthesis (26) and, depending on the sequence of the promoter and the resulting intiation kinetics, may act as either an activator or repressor (31, 36–39). RbpA, on the other hand, was first linked to disulfide stress responses (40) and later found to bind both σ^A^ and σ^B^ holoenzymes, altering transcription initiation kinetics (16, 31, 38, 41). In addition, RbpA and CarD function cooperatively in the context of σ^A^-dependent transcription (31, 33, 37). Given that both regulators affect transcription kinetics rather than directly determining promoter specificity, their combined influence may fine-tune σ factor activity.

Another critical factor influencing transcription in *Mtb* is DNA topology. The energetics of DNA unwinding—modulated by the level of negative superhelicity—affects the efficiency of transcription initiation (42). In bacteria, genomic DNA is typically maintained in an underwound state (43, 44) to facilitate the unwinding reactions necessary for transcription (45–50). Furthermore, genome superhelicity can shift in response to environmental and metabolic cues (51, 52) and, while negative superhelicity generally promotes transcription initiation, its effects can be highly context-dependent (53, 54). For instance, in *E. coli*, relaxed templates enhance σ^38^-dependent transcription and underwound DNA preferentially support σ^70^-dependent transcription (55, 56). Alternatively, in *Bacillus subtilis*, σ^A^-, σ^B^-, and alternative σ factor-dependent transcription appears to be largely driven by negative superhelicity (57). In *Mtb*, it remains unknown whether changes in DNA topology differentially impact σ^A^- and σ^B^-dependent transcription.

This interplay between σ factors, transcription regulators, DNA topology, and promoter architecture is particularly relevant for rRNA transcription, a process that governs protein synthesis and cellular energy balance (58, 59). Since protein translation consumes nearly 80% of a bacterium’s ATP (60, 61), regulating rRNA synthesis is critical for survival, particularly under stress conditions. Unlike many bacteria, which possess multiple rRNA operons (62–64), *Mtb* has only one, known as *rrnA* (65, 66), which can be transcribed from two promoters: the principal *rrn*P3 and the less-characterized *rrn*P1 (66, 67). The extent to which these promoters function under different regulatory contexts remains unclear, particularly regarding σ factor usage and DNA topology. If, as recent data suggest, σ^B^ RNAP holoenzyme transcribes rRNA under specific conditions (9, 25), this would imply an alternative mechanism for maintaining ribosome synthesis during stress or dormancy.

To better understand how σ factor identity, promoter usage, transcription factors like CarD and RbpA, and DNA topology collectively regulate rRNA transcription, we measured steady-state transcription rates across hundreds of reconstituted conditions using a real-time fluorescent-aptamer-based assay (37). This approach allows for direct comparisons of transcription dynamics under different conditions and provides a framework for understanding how these regulatory elements contribute to the *in vivo* regulation of rRNA transcription in *Mtb*. Our findings shed light on the intricate network governing rRNA transcription showing how, under all conditions and factors tested here, the bulk of rRNA synthesis is produced by σ^A^ RNAP holoenzyme on *rrn*AP3.

## RESULTS

### Transcription on the *rrnA* ribosomal promoter is overwhelmingly driven by σ^A^ on the *rrnA*P3 promoter

The mycobacterial rRNA operon belongs to the *rrnA* family which is typically controlled by two or more tandem promoters. For instance, faster-growing species such as *M. smegmatis* possess four promoters that regulate rRNA synthesis, whereas slow-growing species like *Mtb* have only two: *rrnA*P1 and *rrnA*P3 (68). This difference suggests that variations in the utilization of *rrn* promoters could be a strategy for regulating rRNA production. Prior work evaluating the *Mtb* rRNA operon used LacZ reporting plasmids to illustrate that *rrnA*P3 showed consistently stronger signal across all growth stages tested compared to *rrnA*P1 (67). In addition, while no RNA transcripts were detected for *rrnA*P1, the half-life of the *M. bovis* σ^A^ holoenzyme on a combined *rrnA*P13 template was greater than the half-life on *rrnA*P3 alone (69).

Here, we systematically examined the effects of σ^B^ and σ^A^ holoenzymes on the two *rrnA* promoters, *rrnA*P1 and *rrnA*P3 using a fluorescent-aptamer-based steady state transcription assay as previously described (37). We constructed circular DNA templates containing either *rrnA*P1 or *rrnA*P3 individually, or both promoters combined, followed by an iSpinach-D5 aptamer sequence and measured their transcription rates under steady-state conditions with either σ^B^ or σ^A^ RNAP holoenzymes (see Methods). In each reaction, 100 nM purified holoenzyme was pre-incubated with 5 nM *rrnA*P template and transcription was initiated by the addition of 1 mM rNTPs. Transcription was monitored in real time through the fluorescence enhancement produced when the small molecule fluorophore DFHBI binds to the folded RNA aptamer which is transcribed. All experiments were conducted in a 384-well plate-reader format in 10 μl reaction volumes at 37°C. The slope of the fluorescence signal at long times (*i*.*e*., 500 - 1800 s) was used to ensure quantitation of the steady-state rate of transcription without contamination from any initial burst phases (*e*.*g*., as particularly apparent in the σ^B^ traces, Methods). Control experiments using either a template with no promoter (**Figure 1**, dashed black lines, white bar) or core only RNAP (*i*.*e*., no σ factor) **(Figure 1A,C**, dotted black lines) demonstrate that the signals are promoter and holoenzyme specific.

**Figure 1:**
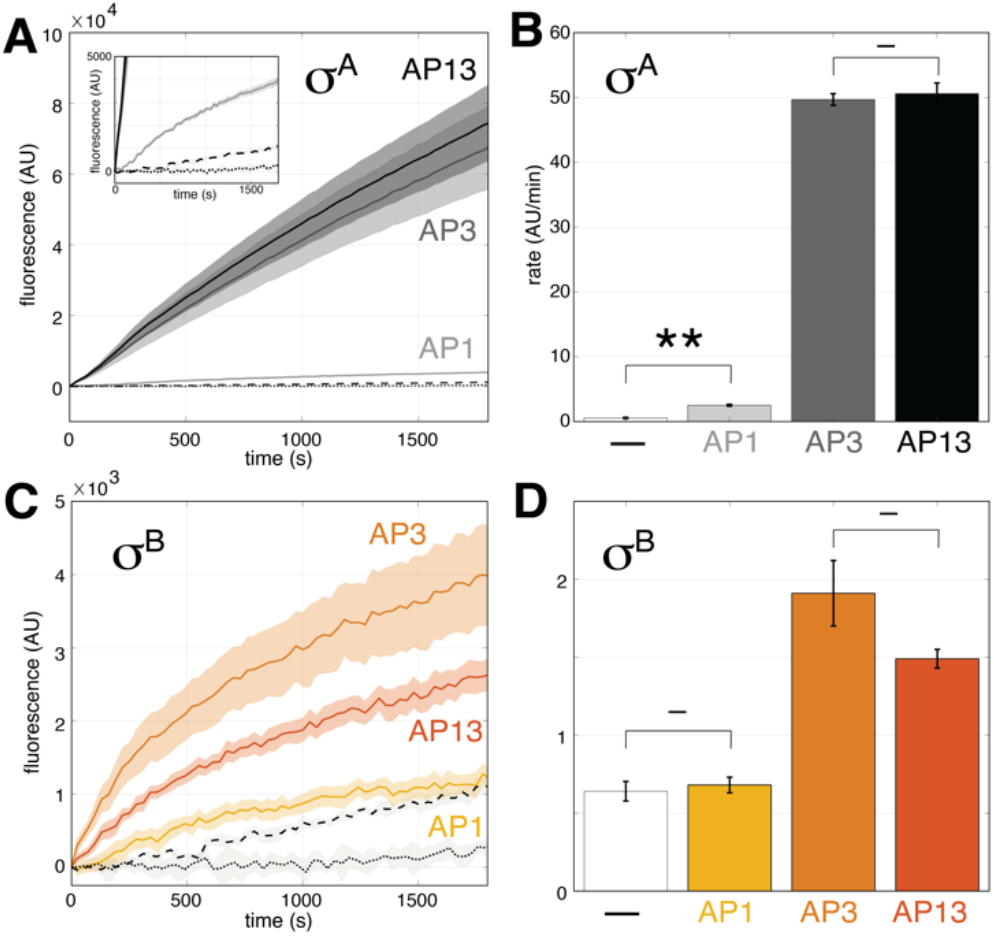
Quantification of steady state-rates on rrnAP1, rrnAP3, rrnAP1AP3, and promoterless circular plasmid templates initiated with rNTPs for σ^A^ (top) and σ^B^ (bottom) holoenzymes. **(A**,**C)** Comparison of real-time fluorescent signal time courses of RNA synthesis. Template without promoter is shown with a black line. Signal from rrnAP3 with core RNAP (i.e., no σ factor) is depicted with a dotted line. Shaded areas indicate the standard error of the mean of 5 experiments. **(B**,**D)** Quantification of steady state rates using linear fits between 500 - 1800 s. Template without promoter is shown in grey. Data are plotted as mean ± SEM of five independent replicates (each one performed in three technical replicates). P-values (paired t-test) are indicated as follows: not significant (–) and less than 0.05 (*). AP3 containing templates are significant over no promoter with P-values of 8×10^−7^ and 2×10^−3^ for σ^A^ and σ^B^ holoenzymes respectively.

The data show that σ^A^ holoenzyme exhibits a 20-fold higher steady-state rate of transcription on *rrnA*P3 compared to *rrnA*P1 (**Figure 1A,B**). In the case of the σ^B^ holoenzyme, this preference for *rrnA*P3 is also observed. *rrnA*P1 transcription by σ^B^ is not statistically different than the template without promoter, while rates from *rrnA*P3 are twice as high. More significantly, overall transcription rates from σ^B^ are over an order of magnitude lower than with the σ^A^ holoenzyme (**Figure 1C,D**).

We also asked whether there may be cooperative effects on the steady-state rate of transcription due to the presence of both promoters, *rrn*AP1 and *rrn*AP3, in tandem. Transcription rates for σ^A^ holoenzymes on the *rrnA*P1P3 construct do not differ significantly from the rates observed for *rrnA*P3 alone (**Figure 1A,B**), and transcription rate averages from the σ^B^ holoenzyme show a small reduction, although the P value indicates no significance (P=0.083) (**Figure 1C,D**). This result indicates that the *rrnA*P1 promoter contributes minimally, if at all, to overall transcription under these conditions, while *rrnA*P3 is predominantly transcribed by the σ^A^ holoenzyme.

### σ^A^ steady-state transcription rate is two orders of magnitude faster than σ^B^ on *rrnA*P3 with similar σ-concentration dependencies

Sigma-factor concentration and availability varies subject to cell growth phases and environmental signals (4, 15, 22, 23, 70–72), causing the concentration of corresponding holoenzymes to change according to the affinity of each σ to core RNAP. To inform on the activities of σ^A^ and σ^B^ holoenzymes on ribosomal transcription, we measured the concentration dependence of both σ factors by titrating untagged σ factors in the multiround transcription assay.

Each reaction contained 100 nM core RNAP, while σ factors concentrations varied between 25 and 1400 nM. Transcription was initiated by adding 5 nM of the *rrnA*P3 circular plasmid template.Due to the initiation of the reaction via the addition of DNA instead of NTPs as above, traces lacked a burst phase (**Supplementary Figure 1**) and were linearly fit to extract steady-state rates. Fits of the titrations with hyperbolic curves (V = V_max_([σ]/(K_m_+[σ])) allowed us to quantitate the concentration at which half activity is reached (K_m_) and the maximal velocity (V_max_). For σ^A^, the K_m_ is 192 ± 50 nM, while for σ^B^, the K_m_ is 177 ± 46 nM (**Figure 2A**). Thus, despite the two-orders of magnitude difference in the maximum velocity at saturating σ factor concentrations (92.5 ± 7.3 and 1.1 ± 0.1 AU/min respectively), σ^A^ and σ^B^ exhibit similar concentration dependencies (**Figure 2A**).

**Figure 2:**
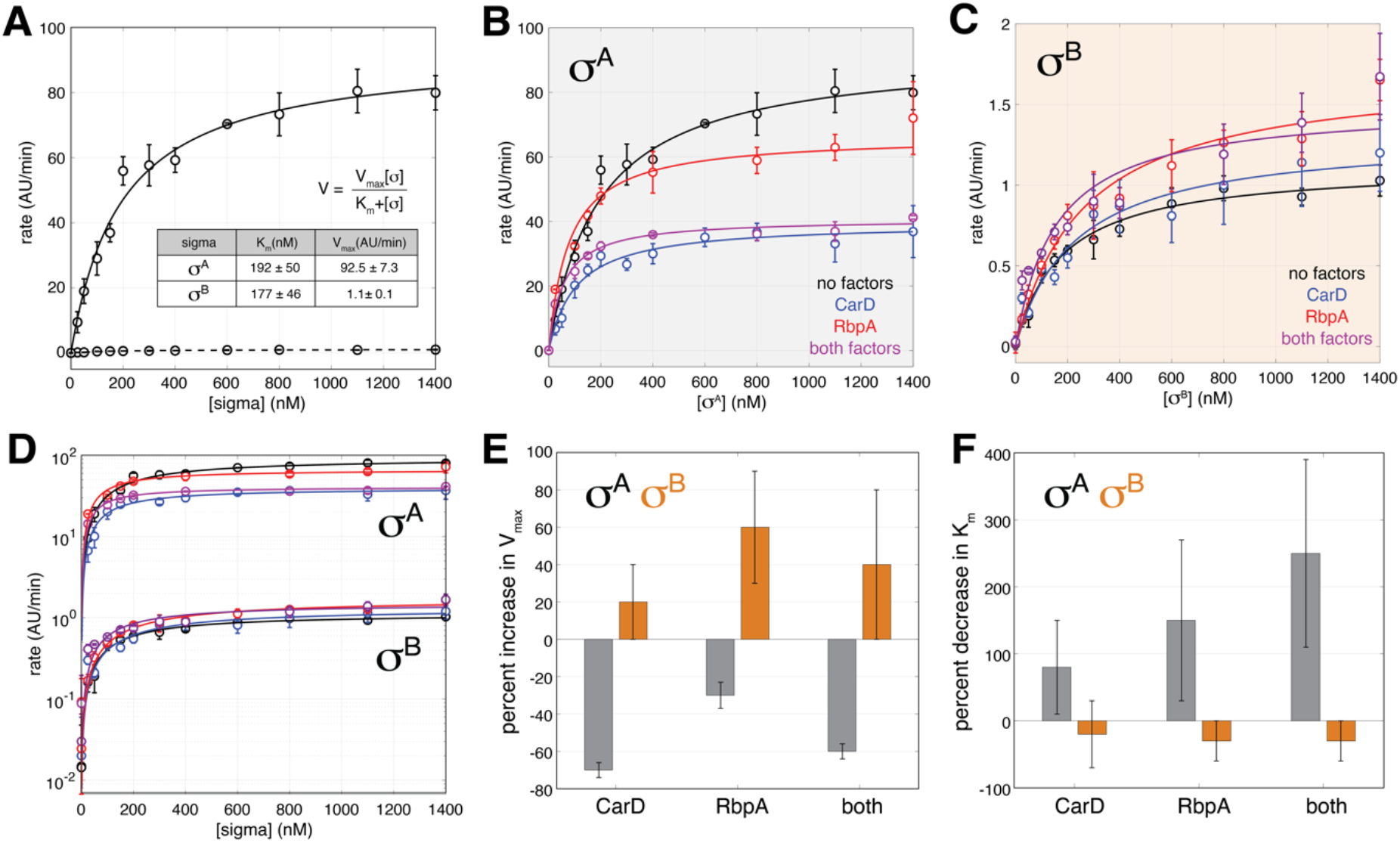
The influence of CarD and RbpA on σ^A^ and σ^B^ driven transcription. **(A)** Steady-state transcription rates as a function of σ factor concentration fit to a hyperbolic curve (σ^A^ solid line, and σ^B^ dotted line). Table insert shows the fit parameters. **(B)** Titration of σ^A^ with no factors (black), CarD (blue), RbpA (red), and both factors (purple). **(C)** Titration of σ^B^ with no factors (black), CarD (blue), RbpA (red), and both factors (purple). **(D)** A comparison of the titrations in B and C on a logarithmic scale. Error bars represent standard error of the mean for each measurement and solid lines represent fits to a hyperbolic curve. **(E)** The percent increase in V_max_ for σ^A^ (gray) and σ^B^ (orange) holoenyzmes calculated from the data in (B) and (C) is plotted for each factor individually and combined. **(F)** The percent decrease in K_m_ for σ^A^ (gray) and σ^B^ (orange) holoenzymes calculated from the data in (B) and (C) is plotted for each factor individually and combined.

### The effects of CarD and RbpA on *rrnA*P3 transcription depend on σ factor identity and are minor compared to the difference between σ^A^ and σ^B^ basal levels

Given the stark decrease in V_max_ with σ^B^ compared to σ^A^ holoenzymes on *rrnA*P3 (**Figure 2A**), we wondered whether the presence of either or both CarD and RbpA may specifically stimulate σ^B^ holoenzymes to the level observed of σ^A^, or if they merely fine-tune the transcriptional output of each. CarD is recruited to initiation complexes in a σ-independent mechanism through interaction with the core RNAP (73). In contrast, RbpA interacts both with core RNAP and with the non-conserved regions of σ^A^ and σ^B^ (16, 33, 35, 41, 74). Given the global nature of these factors (27, 75) their regulation as a function of bacterial growth (26, 76, 77), and their importance in the response to environmental stresses (26, 77, 78), one might hypothesize that these factors are especially important for the regulation of the steady-state rate of ribosomal transcription. To address how transcriptional activity for each holoenzyme varied in the presence of CarD and RbpA, we measured the σ factor concentration-dependence of steady-state transcription on plasmids in the presence of CarD (1 µM), RbpA (2 µM) or both (**Figure 2**). For each σ factor titration, we fit the data to a hyperbolic curve and extracted estimates of the concentration dependence (K_m_) and the maximum velocity (V_max_) (**Figure 2B,C**). We then calculated the ratio of these values to those obtained for each holoenzyme in the absence of factors (**Supplemental Table 1**).

Both factors alone and together repress σ^A^-dependent transcription. Specifically, RpbA reduces V_max_ by 30%, while CarD reduces V_max_ by 70% (**Figure 2B,E**). In contrast, in the case of σ^B^, the factors led to an increase in V_max_ by up to 40-60% depending on the factor (**Figure 2C,E**). Thus, with respect to steady-state transcription on circular plasmid templates, CarD and RbpA have opposite effects on σ^A^ and σ^B^ driven transcription. Furthermore, these effects are small relative to the overall difference between the basal levels of transcription exhibited by each holoenzyme alone (**Figure 2D**). Analysis of the K_m_ values indicates that both factors alone and together decrease the σ^A^-dependence. Specifically, RbpA decreases K_m_ by 80%, while CarD reduces K_m_ by 150% (**Figure 2B,F**). In contrast, in the case of σ^B^, the factors have an distinct effect. Specifically, RbpA increases K_m_ by 30% while CarD did not significantly affect it (**Figure 2C,F**).

### DNA unwinding rate and holoenzyme recycling kinetics limit σ^B^ transcriptional output

In an effort to determine the underlying features that bring about the large changes in V_max_ between the two holoenzymes, we measured aspects of initiation kinetics for both σ^A^ and σ^B^ holoenzymes using linear *rrnA*P3 templates labeled with a Cy3 on the non-template thymine base at position +2 in a stopped-flow assay (31, 36, 79). In the assay, promoter binding and subsequent open complex formation leads to an increase in fluorescence intensity, whereas removal of RNAP from the promoter either via dissociation or promoter escape leads to a decrease.

We first looked at the approach to equilibrium in the forward direction by mixing DNA and RNAP in the absence of NTPs and tracking the fluorescence increase (**Figure 3A**). Based on the time to reach half-maximal signal, σ^A^ holoenzymes equilibrate three times as fast as σ^B^ (t_1/2_s of 31 s and 95 s respectively). Since the equilibration rate reports on the sum of the forward and reverse rates in a two-state system, this observation suggests that either DNA unwinding, DNA bubble collapse, or both are faster for the σ^A^ holoenzyme. Measurements of the dissociation of these complexes by challenging with unlabeled DNA templates provided evidence for an early phase of σ^B^ dissociation that is ∼1.5x faster than that of σ^A^ (**Figure 3B**). Taken together, these observations suggest an ∼4.5x slower rate of DNA opening catalyzed by σ^B^.

**Figure 3:**
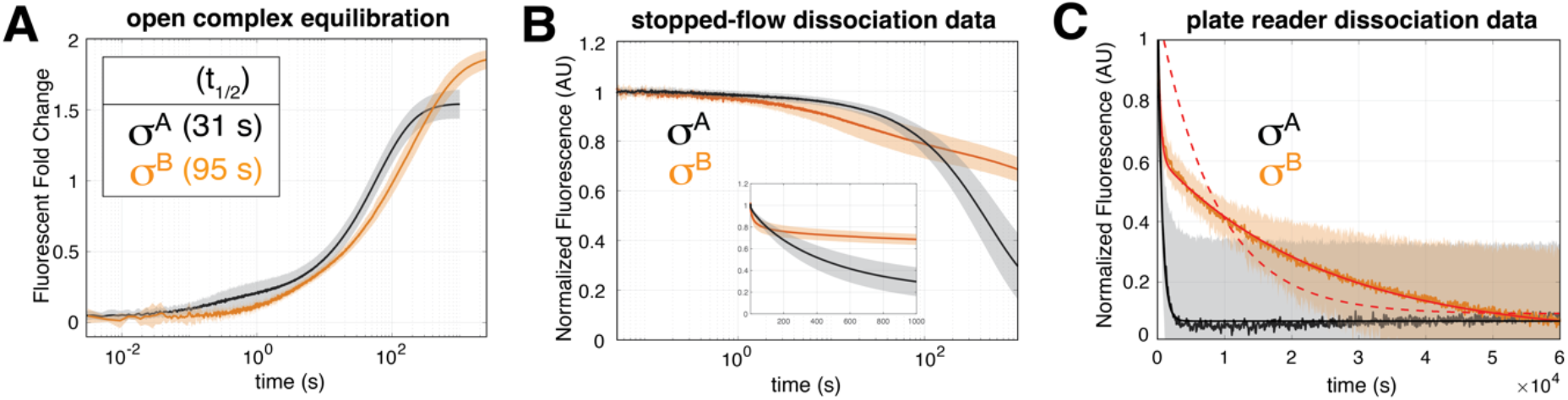
Open complex formation and decay kinetics for σ^A^ and σ^B^ holoenzymes on the rrnAP3 promoter: In all panels, σ^A^ (black) and σ^B^ (orange) holoenzymes are shown. Shaded regions represent the standard deviations from multiple experiments. **(A)** Fluorescent fold-change over background as a function of time showing the approach to open complex equilibrium after the addition of holoenzyme via stopped-flow. **(B)** Open complex decay measured via stopped-flow showing the early phases of dissociation. **(C)** Open complex decay measured via plate-reader showing the longer timescale phase unique to σ^B^ holoenzyme.

We also noted that the σ^B^ holoenzyme saturates at a ∼25% higher fluorescence fold-change (1.85 AU vs. 1.5 AU) suggesting that, under these conditions, σ^B^ eventually forms more open complexes **(Figure 3A)**. This is likely consistent with the presence of an additional and extremely long kinetic phase (t_1/2_ of ∼5 hrs) unique to σ^B^ in the dissociation experiments (**Figure 3B,C**). The multi-phasic dissociation further suggests that σ^B^ populates multiple distinct open complex conformations.

Given the nearly two-orders of magnitude in reduction of the steady-state rate for the σ^B^ holoenzyme, we reasoned that in addition to the ∼4.5x slower DNA unwinding, perhaps promoter escape kinetics would also be reduced. Experiments in the presence of NTPs show that escape for both σ holoenzymes is more rapid than the equilibration of open complex, consistent with the idea that ribosomal promoters are rate limited at DNA opening and open complex formation rather than escape (31, 73). In addition, σ^B^ holoenzyme exhibits a similar rate of decrease in fluorescence compared to σ^A^ (**Figure 4A**), suggesting that promoter escape does not contribute to the relative deficiency of σ^B^-dependent steady-state transcription. Alternatively, we wondered whether the observed NTP-dependent decrease in fluorescence for σ^B^ holoenzyme in this assay could be partially explained via processes that do not lead to the production of a full-length RNA instead of *bona fide* promoter escape. This would reduce the probability of producing a transcript for each open complex formed but would still show a fast decay rate. However, single-round aptamer experiments revealed similar total amounts of transcript produced suggesting that σ^A^ and σ^B^ open complexes have a comparable probability to produce transcript (**Figure 4B**).

**Figure 4:**
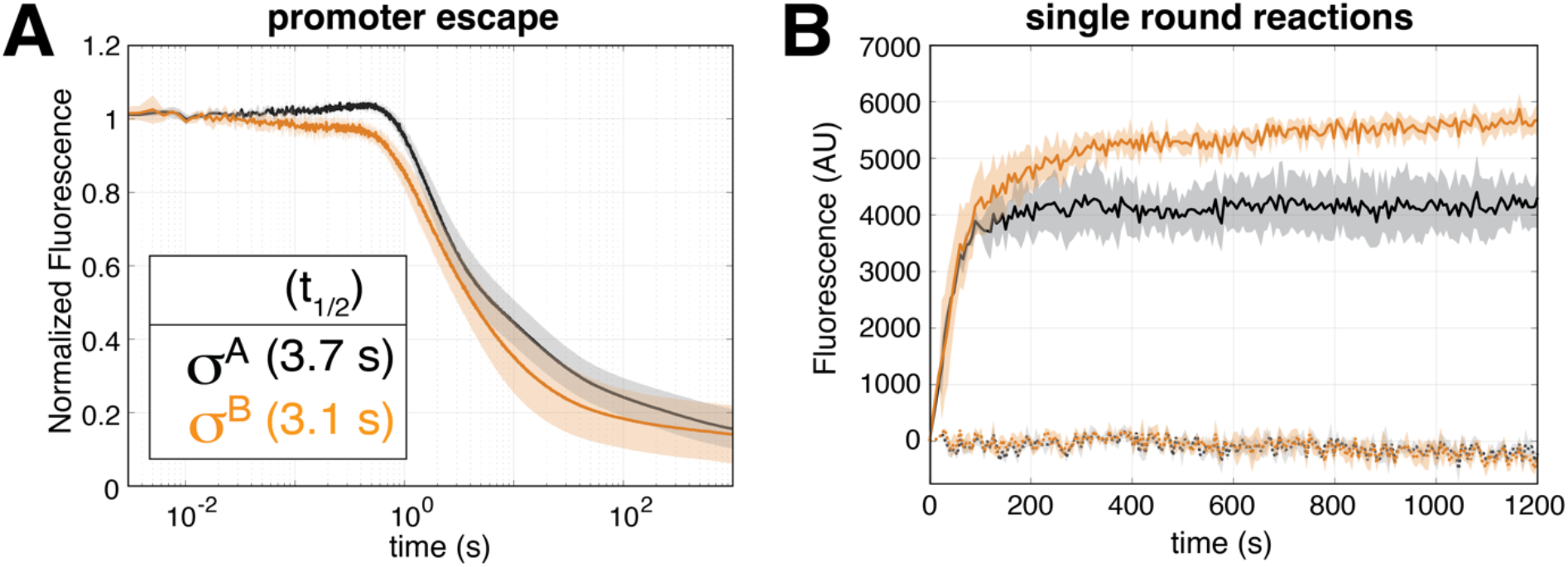
Promoter escape and single round kinetics for σ^A^ and σ^B^ holoenzymes on the rrnAP3 promoter: In all panels, σ^A^ (black) and σ^B^ (orange) holoenzymes are shown. Shaded regions represent the standard deviations from multiple experiments. **(A)** Normalized fluorescence as a function of time showing the decay upon mixing pre-formed open complexes with DNA competitor and rNTPs via stopped-flow. **(B)** Fluorescent signal as a function of time for single-round aptamer assays in the presence of DNA competitor. Both traces were well fit by a single exponential. Controls where DNA competitor was added prior to mixing with NTPs (dotted lines) confirm single-round conditions.

By the process of elimination, these data suggest that in addition to slower open complex formation (**Figure 3A**), the differences in multi-round steady-state transcription must be significantly affected by aspects unique to the multi-round reaction. More specifically, the rates of holoenzyme recycling (*i*.*e*., the rate of σ-core interactions needed to regenerate a competent holoenzyme after transcription) may be a possibility.

### CarD and RbpA produce similar changes in the kinetics of open complex equilibration, dissociation, and promoter escape in the context of σ^A^ and σ^B^ holoenzymes

Above, we showed that σ^A^ and σ^B^ holoenzymes result in different open complex equilibration and dissociation kinetics (**Figure 3**). We also showed that, under the conditions tested, on circular plasmid templates CarD and RbpA repress steady-state transcription by σ^A^ and activate steady-state transcription driven by σ^B^ holoenzymes (**Figure 2**). From theoretical (31, 80, 81) and experimental (39, 82, 83) perspectives exploring how different basal kinetics can result in differential CarD and RbpA-based transcriptional regulation, we hypothesized that CarD and RbpA modulate the kinetics of σ^A^ and σ^B^ holoenzymes in a similar way even though they result in diametrical effects on steady-state rate (**Figure 2**). If this hypothesis is true, each factor would be predicted to accelerate the equilibration of open complexes and slow promoter escape (31) regardless of σ factor context. This prediction is based on the following prior work: i) on *Mtb rrnA*P3 with σ^A^ holoenzymes, CarD and RbpA increase the rate of DNA opening (31, 33, 38) and slow escape kinetics (31), and ii) on *Mtb sigA*P with σ^B^ holoenzymes, RbpA facilitates open complex formation (16, 35, 83). Alternatively, we wondered if, as has been suggested for RbpA (16, 41), there are σ-dependent differences in the activity of each transcription factor. If this picture were true, one might expect that CarD and RbpA would produce different kinetic changes on the same step of intiation depending on σ context.

To test these two possibilites, we performed our kinetic stopped flow assays for each σ factor-containing holoenzyme in the presence of CarD and RbpA individually and combined (**Figure 5**). In each of the two kinetic assays, we observed that the factors changed the kinetics in the same direction. Regardless of σ factor identity, CarD and RbpA accelerated open complex equilibrium (**Figure 5A,B**) and slowed promoter escape (**Figure 5C,D**) and open complex dissociation (**Supplementary Figure 2**). This result suggests that the distinct effects of a particular factor on steady-state transcription rate are dictated by the different basal kinetics of σ^A^ and σ^B^ and not by distinct molecular mechanisms. That said, while the magnitude of change in the kinetics due to CarD is similar in the two σ factor contexts (**Supplementary Figure 3A**), RbpA exhibits a quantitatively stronger effect on the kinetics of σ^B^ (**Supplementary Figure 3B**).

**Figure 5:**
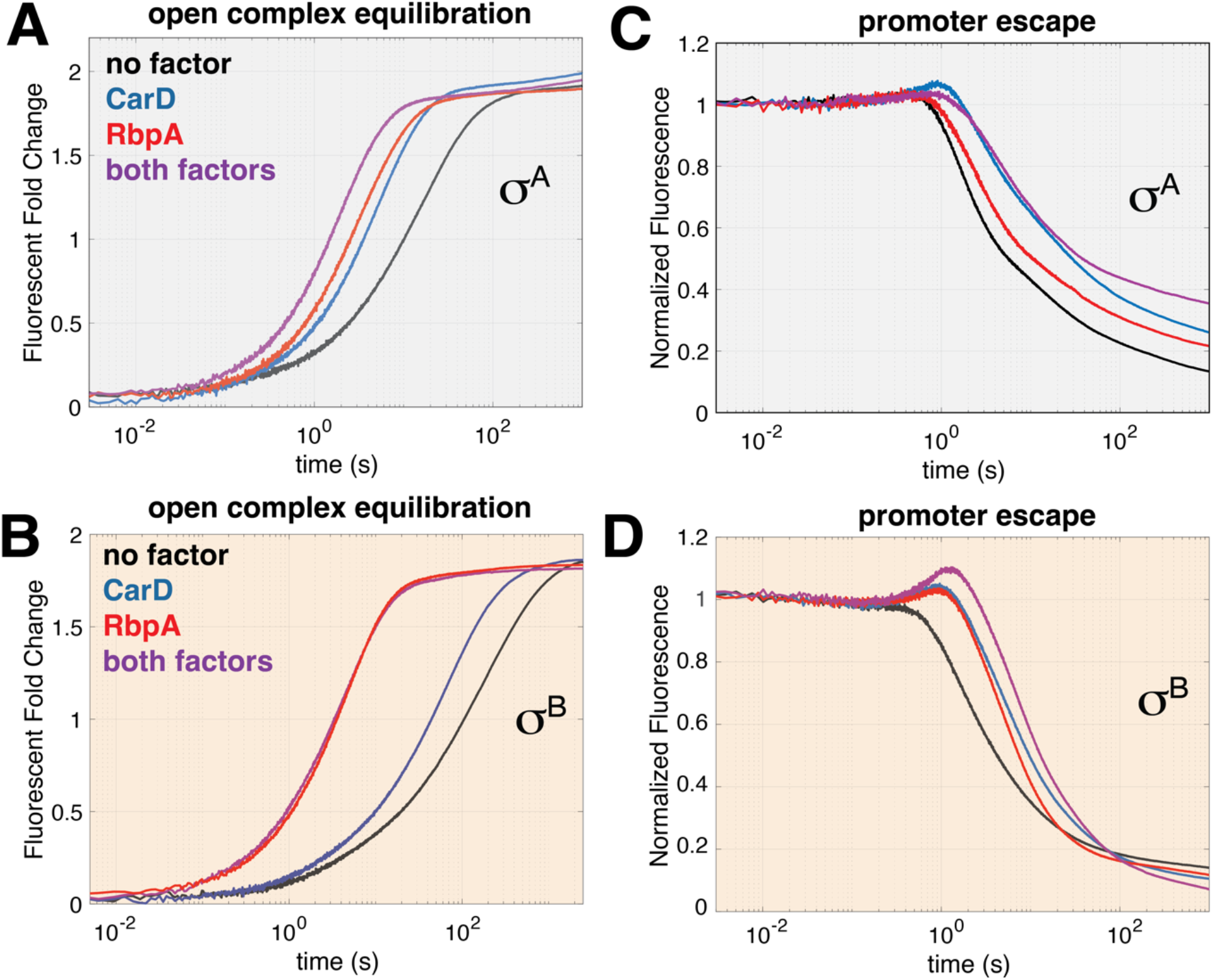
Dependence of forward and dissociation/escape kinetics on CarD and RbpA for σ^A^ and σ^B^ holoenzymes on the rrnAP3 promoter: In all panels, no factor (black) traces were collected under conditions where open complex was saturated and are compared to those in the presence of CarD (blue), RbpA (red), and both factors together (purple). **(A**,**B)** Open complex equilibration kinetics for σ^A^ and σ^B^ holoenzymes respectively. **(C**,**D)** promoter escape kinetics for σ^A^ and σ^B^ holoenzymes respectively. Note that the increase in fluorescence starting at ∼ 100 ms before the fluorescence decay reports on initial transcribing intermediates (see (31)) and is not discussed here further.

### Negative superhelicity stimulates σ^A^ and σ^B^ driven *rrnA*P3 transcription equally

Knowing that DNA topology dramatically affects the energy landscape of DNA unwinding (42), we next asked how steady-state transcription driven by each σ factor-containing holoenzyme, and its transcription factor-dependence, varies with templates exhibiting differing levels of superhelical densities. To investigate the influence of DNA topology on transcription rates, we utilized a set of five *rrnA*P3 DNA templates (**Supplementary Figure 4** and Methods): supercoiled (miniprepped and DNA gyrase treated), relaxed (Topoisomerase I treated), nicked (Nt.BsmAI treated), and linearized (ScaI treated). Gyrase treated templates showed comparable rates to the miniprepped DNA and are not discussed further (**Supplementary Figure 5**).

We first measured basal rates with either σ^A^ or σ^B^ holoenzymes in each topology. In both holoenzymes, the supercoiled template stimulated steady-state transcription by 2-3-fold compared to nicked and linear templates with the relaxed template displaying an intermediate rate (**Figure 6, gray**). Overall, the ratios between σ^A^ and σ^B^ transcription rates are relatively constant across the different topologies (**Supplementary Table 3**, no factor columns).

**Figure 6:**
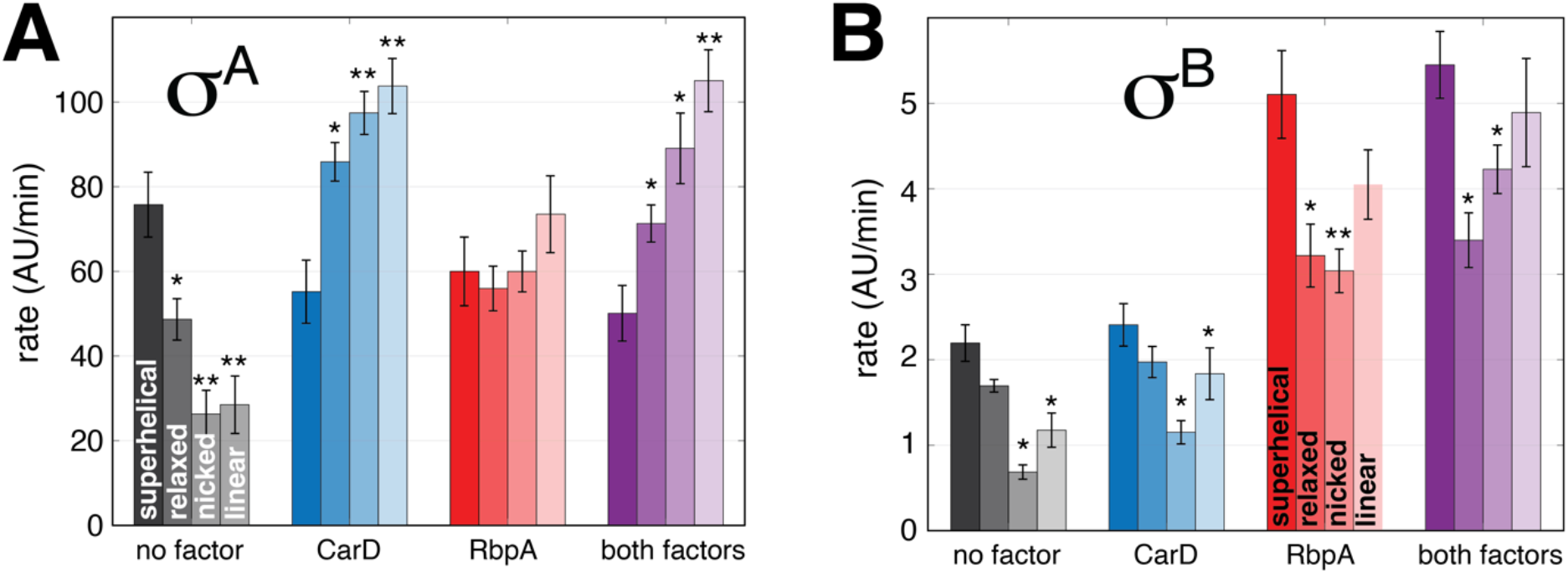
The dependence of steady-state transcription on topology and transcription factor for σ^A^ **(A)** and σ^B^ **(B)**. Factor conditions are indicated on the x-axis and topology is indicated by different shades. Error bars indicate standard error of the mean from at least 6 traces in each condition collected across multiple days, each with triplicate technical replicates. P-values (paired t-test) from the supercoiled template are indicated by asterisks as follows: less than 0.05 (*), less than 0.005 (**), less than 0.0005 (***).

### CarD’s and RbpA’s effects on steady-state rate differ with respect to topology and σ factor identity

#### CarD and RbpA reduce or invert the topology dependence of σ^A^ transcription on rrnAP3

In the presence of CarD, and agreeing with previous observations (39), the rate of σ^A^-dependent transcription on the superhelical template is slightly reduced while that of relaxed, nicked, and linear are significantly enhanced (**Figure 6A**, grey vs. blue). In fact, the magnitudes of the changes are such that the topology dependence is completely reversed in the presence of CarD. RbpA also slightly reduced the transcription rate on the supercoiled template and stimulated transcription on the other templates such that any topology dependence was nearly completely removed (**Figure 6A**, grey vs. red). In the presence of both transcription factors, the reversed topology dependence driven by CarD is again visible (**Figure 6A**, purple). The same data are grouped in terms of topology for a comparison of the effect of each transcription factor on a given template (**Supplementary Figure 6A**). Here one can more clearly see that CarD increases rates of transcription more than RbpA in the context of σ^A^. The fold changes relative to no factors and relative to the supercoiled templates can be found in **Supplementary Tables 2 and 3**.

#### RbpA activates transcription across all topologies in the context of σ^B^

CarD has a limited effect on the transcription rates of σ^B^ holoenzyme (**Figure 6B**, gray vs. blue and **Supplementary Figure 6B**), consistent with limited effects on open complex equilibration and promoter escape (**Figure 5**). In contrast, RbpA significantly stimulates transcription on all topologies (**Figure 6B**, gray vs. red and **Supplementary Figure 6B**), also consistent with displaying larger effects than CarD on individual initiation steps (**Figure 5**). In the presence of both transcription factors, the effect is similar to that of RbpA alone (**Figure 6B**, purple vs. red) As such, the preference for supercoiled templates is conserved across all conditions. That said, the stimulator effect of RbpA is greater on the nicked and linear templates compared to the supercoiled template (2x vs. 4x) and less so on the relaxed template (1.5x) (**Supplementary Figure 6B**). Calculated fold changes relative to no factors and relative to the supercoiled templates can be found in **Supplementary Tables 2 and 3**.

### Differences in steady-state transcription rate are linked to the N-terminal extension of σ^A^

Given the stark disparity in rRNA transcription rates between σ^B^ and σ^A^ holoenzymes in all conditions tested here, we considered potential structural determinants behind this difference. The most notable distinction between the two σ factors is the presence of a 205 residue N-terminal extension in σ^A^ that is almost certainly intrinsically disordered (74). Although the exact mechanism by which this feature modulates σ^A^ function remains unknown, single timepoint gel-based experiments on linear *rrnA*P3 templates suggest that, while deleting the first 179 residues of the N-terminal extension may or may not affect promoter binding, transcriptional output is reduced four-fold relative to full-length σ^A^ (9). Additionally, this truncation failed to complement an *in vivo* model suggesting an essential functional role (9). Thus, we tested whether the N-terminal extension in σ^A^ accounts for the observed differences between σ factors in the steady-state rates of r*rnA*P3-driven transcription.

Using a mutant version of σ^A^ (σ^A^_Δ205_) in which the first 205 amino acids were deleted, we performed steady-state transcription reactions as a function of σ concentration. Each reaction contained 100 nM core RNAP while the concentration of the mutant σ factor was titrated from 25 to 1100 nM (**Figure 7A**). The data show that σ^A^_Δ205_ has a V_max_ two orders of magnitude lower than wildtype σ^A^ and similar to σ^B^ (**Figure 7A,B**). Thus, the N-terminal extension directly contributes to the high steady-state rates observed with σ^A^. We also tested the effect of transcription factors CarD and RbpA on σ^A^_Δ205_. These results suggest that the σ^A^ N-terminal extension also dictates the factor-dependent effects (**Figure 7B, Supplementary Table 1**).

**Figure 7:**
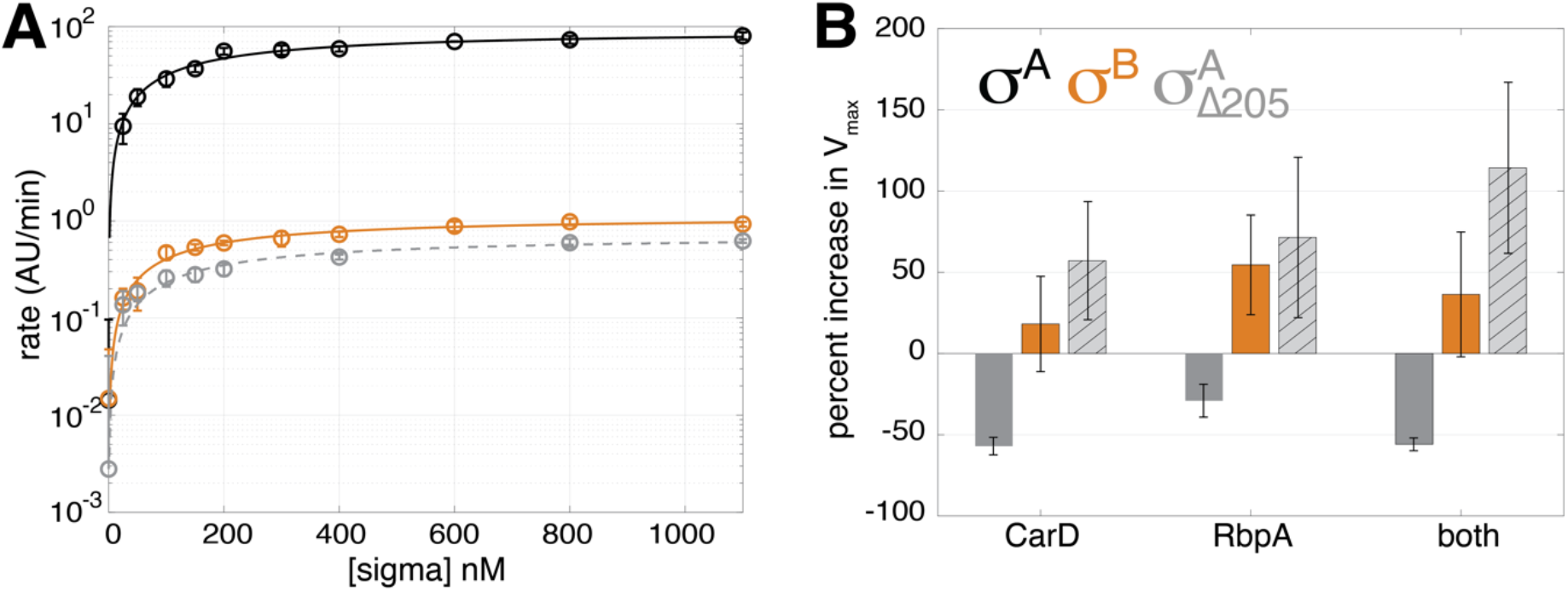
The effecct of the N-terminal extension of σ^**A**^. **(A)** Steady state transcription rates as a function of σ factor concentration for σ^A^ (black), σ^B^ (orange), and σ^A^_Δ205_ (dashed gray line). Error bars represent standard error of the mean and lines represent fits to hyperbolic curves. (**B)** Percent changes in V_max_ for each σ factor (σ^A^:dark grey, σ^B^:orange, σ^A^_Δ205_:light gray with hatchmarks) in the presence of each transcription factor.

## DISCUSSION

The expression of ribosomal rRNA genes is tightly regulated in response to environmental signals. Both the housekeeping sigma factor A (σ^A^) and the alternative sigma factor B (σ^B^) are expressed at roughly equal levels during exponential growth (23) and are associated with similar promoters, including rRNA promoters, as revealed by ChIP-seq (25). While the σ^B^ regulon overlaps significantly with that of σ^A^, and it is classified as a secondary σ factor that may complement σ^A^ function during growth (9, 25, 84), it is not essential and cannot rescue a σ^A^ knockout (9). Notably during hypoxia or stationary phase, conditions that contribute to the transition of *Mtb* into a non-replicative, dormant state, σ^A^ expression decreases 3-fold while σ^B^ levels increase 2.5-fold (23). This coincides with a reduction in ribosomal synthesis (85), albeit to a lesser extent compared to other bacteria (86). Here, we investigated the degree to which rRNA transcription by σ^B^ holoenzymes contribute to the regulation of rRNA synthesis.

### Ribosomal transcription output is dominated by σ^A^ holoenzymes from the rrnAP3 promoter

Our steady-state measurements show that *rrnA*P3 exhibits dramatically higher output than *rrnA*P1 and that the presence of both promoters does not significantly alter the rate of transcription (**Figure 1**). In addition, σ^B^ holoenzymes exhibit a notably lower output compared to σ^A^ holoenzymes across all three promoter constructs (*rrnA*P1, *rrnAP1P3*, and *rrnA*P3) (**Figure 1**). However, these initial measurements only included the basal transcription machinery and lacked CarD and RbpA, two global transcription factors that are often considered to be *de facto* members of the complex as they are each recruited directly to initiation complexes by the RNAP holoenzyme itself (33, 73). We considered the possibility that the presence of these transcription factors may dramatically change the picture of how much σ^B^ contributes to ribosomal transcriptional output. However, while each transcription factor clearly regulates steady-state transcription driven by each holoenzyme, factor-dependent rate changes are small relative to the nearly two orders of magnitude difference between the rates of σ^A^ and σ^B^ (**Figure 2**).

Taken together, our results suggest that despite observations of σ^B^ holoenzymes at ribosomal promoters (25) and the downregulation of σ^A^ under stress and in stationary phase (23), rRNA transcription is likely to be primarily driven by σ^A^ holoenzymes at the *rrnA*P3 promoter. This result underscores the central role of σ^A^ and provides constraints when considering the role of σ^B^ in maintaining ribosomal transcription. Yet-to-be-discovered regulatory mechanisms enacted under specific growth conditions in specific environments *in vivo* may change this picture.

### Mechanisms underlying differential ribosomal transcription of σ^A^ and σ^B^ holoenzymes

Our kinetic measurements revealed that σ^B^ holoenzymes exhibit distinct initiation kinetics in that they have a slower rate of open complex equilibration, a very long-lived open complex in the absence of NTPs, and a similar promoter escape rate compared to σ^A^ holoenzymes (**Figures 3**,**4**). However, these kinetic and structural studies appear to represent only part of the story with respect to large differences in steady-state rates. Since each holoenzyme exhibits similar single-round kinetics (**Figure 4B**), we propose that differences in the rate of holoenzyme recycling (*i*.*e*., the re-binding of σ and core after a round of transcription) may impact the differences in steady-state rates (discussed further in the section below devoted to the role of the N-terminal extension). (87–89)(89)

We also note work showing that N terminally tagged σ^B^ holoenzymes form higher order oligomers that may be inhibitory to transcription (35). These complexes begin to form at concentrations of holoenzyme used here. However as they are dissolved by RbpA and we show that the presence of RbpA does not equalize the rates between σ^A^ and σ^B^ holoenzymes (**Figure 2D**), the effect of these oligomers is likely subtle. Taken together, it appears that multiple mechanisms combine to determine the relative rates of ribosomal transcription by the different holoenzymes.

### CarD and RbpA have opposite effects on the rate of transcription of σ^A^ and σ^B^ holoenzymes on negatively supercoiled DNA

In the context of superhelical templates, the presence of CarD and RbpA exert opposite effects on σ^A^ and σ^B^ driven transcription. Specifically, they repress *rrnA*P3 transcription mediated by σ^A^ while activating transcription mediated by σ^B^ (**Figure 2**). These opposing effects on the maximal steady-state rate are paralleled by changes in the concentration dependence of each σ factor: CarD and RbpA reduce the K_m_ for σ^A^, consistent with previous studies with RbpA (41), while increase it for σ^B^, with an approximate two-fold difference in each case (**Figure 2E**,**F and Supplementary Table 1**). To explore the mechanistic basis of these trends, we considered two potential explanations. The first is that a transcription factor can lead to different regulatory outcomes depending on the basal kinetics of a given promoter/holoenzyme system (31, 80). The second is that a transcription factors exhibits σ factor specificity, altering initiation kinetics in a σ-dependent manner to either repress or activate transcription (16, 41).

To distinguish between these models, we examined the effects of the CarD and RbpA on three key aspects of initiation kinetics using a fluorescence stopped-flow assay as previously described (31, 38, 87). Our results confirmed that CarD and RbpA induce qualitatively similar changes in the kinetics of open complex equilibration, open complex dissociation, and promoter escape, regardless of σ factor identity (**Figures 3, 4, Supplementary Figure 2**). However, within this context, RbpA shows a quantitatively enhanced ability to modulate these kinetics in the presence of σ^B^ compared to σ^A^ (**Supplemental Figure 3**). The ability of RbpA to dissolve inactive σ^B^ holoenzyme octamers could contribute to this specificity (35).

Thus, we are left with a mixture of the two models. Overall, the direction of the regulatory outcome arises from a shared transcription factor mechanism modulated by the basal initiation kinetics of each promoter/holoenzyme system. Quantitatively, there also appears to be molecular specificity, with RbpA showing a preference for enhancing σ^B^ holoenzyme activity. Together, these results point to a complex interplay of basal kinetics and σ−specific interactions in determining transcriptional regulation by CarD and RbpA.

### Negative superhelicity stimulates ribosomal transcription by both σ^A^ and σ^B^ holoenzymes

Negatively supercoiled DNA is maintained during growth by a balance between ATP-dependent DNA gyrase and other topoisomerases (88). As such, genome superhelicity is also influenced by cellular energy stores (*i*.*e*., ATP/ADP ratios). For example, during stationary phase, when energy becomes limited and ATP levels drop, reduced gyrase activity leads to decreased superhelicity (89). Conversely, environmental stresses like osmotic or cold stress have been shown to increase DNA supercoiling (hypercoiling) in *E. coli* and *B. subtilis*, which can activate transcription (88). Similarly, in *Mtb*, superhelicity has been implicated in the regulation of virulence genes, such as *virR* and *sodC*, through the *NapA* nucleoid-associated protein (90).

Given the influence of supercoiling on transcription, we investigated how superhelical density affects transcription driven by each σ factor. Consistent with early *in vitro* studies using isolated *M. smegmatis* RNAP (91), we observed that both σ^A^ and σ^B^ driven transcription is strongly dependent on the superhelical state of the DNA template (**Figure 6**). Specifically, transcriptional output by both σ factor-containing holoenzymes is reduced by approximately 50% when comparing supercoiled plasmid templates to linear templates. As negative superhelicity promotes DNA unwinding (45–49), these observations are also consistent with a model where *rrnA*P3 is rate limited in part by DNA opening (**Figures 3A**,**4A**, (31)).

### CarD and RbpA buffer the dependency of σ^A^ holoenzymes on superhelicity

To understand how the intersection of superhelicity and transcription factors influence rRNA transcription, we examined the effects of CarD and RbpA across different σ factor and superhelical contexts. Remarkably, we found that each factor individually, or both factors combined, generated a flip in the dependency of σ^A^ holoenzymes on superhelicity. Specifically, in the presence of both factors, steady-state transcriptional output is stimulated up to two-fold on relaxed, nicked, and linear templates compared to supercoiled plasmid templates (**Figure 6**). This result is consistent with prior work illustrating that with the σ^A^ holoenzyme, both CarD and RbpA activate *rrnA*P3 transcription on linear templates (37, 39, 41, 92), whereas CarD represses transcription on supercoiled plasmid templates (39). This topological dependence on CarD activity may also explain why *in vivo*, CarD depletion in *M. smegmatis* leads to increased rRNA levels (26), assuming CarD directly represses rRNA transctiption on the supercoiled genome. As both DNA topology and promoter sequence may moduulate basal transcription initiation kinetics, our observations are consistent with prior work illustrating how changes in promoter sequence can lead to differential regulation by *Mtb* CarD and RbpA (39, 83) and *Rhodobacter sphaeroides* CarD (82, 93).

In the case of σ^B^ holoenzyme, a similar effect is observed, where CarD and RbpA change the relative output. However in this instance, they reduce the disparity between linear and supercoiled template outputs, bringing them to roughly equal levels **(Figure 6)**. This behavior suggests that CarD and RbpA help buffer reductions in rRNA transcription during stationary phase, when superhelicity and σ^A^ levels are lower (23, 41, 88, 94).

On a practical experimental level, we emphasize the need to carefully control for DNA template topology in *in vitro* studies. Differences in topology, whether due to the use of linear versus circular templates or to the degradation of superhelicity via nicks in plasmids, can significantly affect the results of transcriptional measurements and the inferred biological mechanisms of gene regulation.

### The N-terminal extension of σ^A^ is a major regulatory determinant of the steady-state transcription rate differences between σ^A^ and σ^B^ and the effects of CarD and RbpA

Finally, to explore potential structural determinants of the observed differences between σ^A^ and σ^B^, we examined the unique N-terminal domain extension in σ^A^. Structural studies have suggested that the N-terminal extension of σ^A^, which σ^B^ lacks, likely plays a role in open complex formation/stabilization due to its position in relation to the downstream DNA channel (34, 35, 95, 96). In *Mtb*, this region is predicted to be entirely disordered (74), while the non-homologous N-terminal domain in *E. coli* σ^70^, termed region 1.1, is partly structured and conformationally dynamic (97, 98). For instance, in its apo-form, σ^70^_1.1_ prevents DNA binding by occluding the DNA binding domains in a compact structure (99–101). While bound to core RNAP in a more expanded conformation, σ^70^_1.1_ occupies the active site cleft where DNA gets loaded and must be displaced upon DNA unwinding (97, 98, 102), likely resulting in the observed changes in unwinding kinetics (103) and complex stability (104, 105). Given the lack of sequence and structural conservation between the *Mtb* σ^A^ N-terminal extension and *E. coli* σ ^70^_1.1_ (74), the function of the σ^A^ N-terminal extension remains unknown. As our kinetic measurements suggest that part of the difference in steady-state rates may be due to differences in holoenzyme recycling rates, it is possible that the N-terminal extension also affects this process.

Under the conditions explored in this paper, truncation of the first 205 amino acid residues of the σ^A^ N-terminal extension (σ^A^_Δ205_) generates steady-state rates of transcription comparable to those of σ^B^ on plasmid templates (**Figure 7A**). As such, we conclude that the N-terminal extension is a major determinant of the increased rate of transcription catalyzed by σ^A^ holoenzymes relative to σ^B^. Interestingly, while large changes in V_max_ were observed (**Figure 7B**), the EC_50_ of σ^A^_Δ205_ was not significantly different from the EC_50_ of either σ^B^ or σ^A^, suggesting a similar affinity of each holoenzyme to promoter DNA (**Supplementary Table 1**). It is worth noting that the regulatory effect of CarD and RbpA is greatly influenced by the N-terminal extension, leading to repression of transcription when present (σ^A^) and activation of transcription when absent (σ^A^_Δ205_), analogous to the effects observed with σ^B^ (**Figure 7B**). As a result, CarD and RbpA can be added to the list of other general bacterial transcription factors, like *E. coli* DksA and TraR (102, 104, 106), and bacteriophage proteins (107, 108) whose activity is impacted by the N-terminal region of σfactors.

Given the dramatic effect of the N-terminal extension on basal and regulated kinetics measured here, we stress that σ factor comparisons made throughout this work utilized untagged proteins to avoid potential artefacts of non-native sequences. Clearly, the N-terminus has the potential to alter the complex energetics and kinetics of interactions between σ factors, the RNAP core, and the DNA template.

It is tempting to hypothesize about the regulatory advantage provided by the N-terminal extension into *Mtb’*s ability to balance transcriptional efficiency with adaptability across different environmental contexts. While the exact molecular mechanisms remain to be elucidated, possibilities include modulation of interactions with core RNAP subunits and sequence-specific lineage insertions (74, 109), or other regulatory proteins (110, 111). Future studies will be needed to determine more specific mechanisms underlying σ^A^’s enhanced output to offer insights into the N-terminal extension’s role in the biology of *Mtb*.

In conclusion, ribosomal transcription in *Mtb* is primarily driven by σ^A^, with σ^B^ playing a more limited role under the conditions studied. CarD and RbpA reverse the dependency of σ^A^ holoenzyme on superhelicity (and buffer that of σ^B^ holoenzyme) and have opposing effects on the transcription rates of σ^A^ and σ^B^ holoenzymes on negatively supercoiled DNA. These findings highlight the central role of σ^A^ in meeting *Mtb*’s ribosomal demands and underscore the importance of integrating direct measurements of steady-state transcription rates with techniques like ChIP-seq and RNA-seq for a more comprehensive understanding of the regulation of gene transcription. Further investigation into the interplay between σ factors, DNA topology, and regulatory proteins will provide critical insights into *Mtb*’s adaptability and survival strategies.

## MATERIALS AND METHODS

### Preparation of DNA constructs

For all final DNA construct sequences used in this work, see **Supplementary Table 4**. Circular plasmid templates, 2557 base-pair (bp) in length, were ordered from Twist Bioscience (San Francisco, CA) and represent an updated version of previously published constructs (37). Sequences contain, in order, the *Mtb tuf* terminator (112), the *Mtb* ribosomal RNA promoter (*rrnA*P3), the iSpinach D5 aptamer (113), and the *E. coli rrnB*P1 *T*_1_ terminator (114, 115).

For the preparation of topologically different DNAs, the plasmid purified from the Qiagen Maxi Prep Kit (12963) was more than 90% supercoiled, estimated by the relative abundance of the bands on the agarose gel (**Supplementary Figure 4**). For additional negative supercoiling and removal of concatemers, the plasmid was treated with DNA Gyrase (TopoGEN, TG2000G-1). Additionally, topologically closed but relaxed plasmids were prepared by incubation with Topoisomerase I (NEB, M0301S). Nicked plasmid DNA was prepared by incubation with restriction enzyme Nt.BsmAI (NEB, R0121S) that made two cuts in the plasmid upstream the promoter region. To linearize the plasmid, the restriction endonuclease enzyme ScaI (NEB, R3122S) was used. In all cases, a total volume of 200 μl, 20 μl assay rCutSmart buffer (NEB) 150 μl RNase-free water, 10 μl enzyme and 5 μl of plasmid DNA (c = 2000 ng/μl) were combined in a reaction tube and incubated for 1 h at 37°C. Reactions were cleaned with a PCR clean-up kit (Qiagen, 28104), and resuspended to 50 nM in transcription buffer (see below). Concentrations were calculated using the molecular weight based on sequence.

A Linear 150 bp template containing the *Mtb rrnA*P3 promoter labelled with Cy3-NHS (Lumiprobe Corporation, 11020) attached to a C6-amine modified thymine on the +2 position of the non-template strand was used for plate-reader dissociation and stopped-flow assays. A biotin molecule was added to the 5’-end of the non-template strand attached via a standard C6 spacer (IDT). For promoter preparation and labeling protocols, see (36).

### Preparation of recombinant proteins

A summary of all expression constructs used can be found on the **Supplementary Table 5**.

#### Mtb RNAP core for in vitro reconstituted holoenzymes

To obtain the *Mtb* RNAP core complex (β, β’, His-tagged α, and *ω*), *E. coli* BL21(DE3) cells were transformed with plasmids pET-Duet-*rpoB*-*rpoC* (encoding the β and β’ subunits) and pAcYc-His*rpoA*-*rpoZ* (encoding the N-terminal 10X His-tagged α and *ω* subunits) and were grown in LB media at 37°C to an OD_600_ of 0.6–0.8. Protein expression was then induced with 0.25 mM IPTG and grown overnight at 16°C. Cells pellets resuspended in 20 mM Tris, pH 8.0, 5 mM Imidazole, 500 mM NaCl, 5 mM β-ME with 1 protease inhibitor cocktail tablet (Roche, 05892791001) and lysed via sonication. Clarified cell lysate (via centrifugation at 5,000 rpm for 30 min at 8°C) was loaded on two 5 ml Ni^2+^ HisTrap FF crude affinity columns (Cytiva, 17528601) set up in tandem using a 5–1000 mM imidazole gradient in 20 mM Tris, pH 8.0 and 500 mM NaCl. Fractions of interest were dialyzed overnight (25 kDa MWCO tubing, Spectrum Labs, 132554) in 10 mM Tris, pH 8.0, 250 mM NaCl, 0.1 mM EDTA, 1 mM MgCl_2_, 10 μM ZnCl_2_, and 10 mM DTT, concentrated, and further purified by size exclusion chromatography (HiPrep 16/60 Sephacryl S-300 column, Cytiva, 17116701). Eluted complexes were dialyzed overnight in 10 mM Tris, pH 8.0, 200 mM NaCl, 0.1 mM EDTA, 1 mM MgCl_2_, 20 μM ZnCl_2_, 50% (v/v) glycerol, and 2 mM DTT, defined here as storage buffer. *Mtb* RNAP core was concentrated to ∼5 μM (100 kDa MWCO Vivaspin 20 filters, Sartorius, VS2041), flash frozen in liquid nitrogen and stored at –80°C. Concentration was determined with A_280_ with an extinction coefficient of 245,000 M^-1^ cm^-1^.

#### Co-expressed/purified *Mtb* σ^A^ *RNAP holoenzyme*

Co-expression/purification of the *Mtb* RNAP σ^A^ holoenzyme complex was obtained in the following two ways: 1) use of a 10X N-terminal His-tag on α using pET-Duet-*rpoB*-*rpoC*, pAcYc-His*rpoA*-*rpoZ*, and pAC27-*sigA* plasmids and 2) use of a 10X N-terminal His-tag on σ^A^ using pET-Duet-*rpoB*-*rpoC*, pAcYc-His*sigA*-*rpoA*, and pCDF-*rpoZ* plasmids (used only in 2 experiments in the comparison of *rrnA* promoters section with no difference in activity). Complexes were expressed and purified as described for *Mtb* RNAP core above. In some cases, an anion exchange chromatography step (MonoQ 10/100 GL, Cytiva, 17516701) was added. Fractions of interest were dialyzed overnight into 20 mM Tris, pH 8.0, 150 mM NaCl, 0.5 mM EDTA, 1 mM MgCl_2_, 5% (v/v) glycerol and 2 mM β-ME and purified using 150–1000 mM NaCl gradient. The final holoenzyme fractions were dialyzed into storage buffer, concentrated and frozen as described for *Mtb* RNAP core, using an extinction coefficient of 280,425 M^-1^ cm^-1^.

#### Co-expressed/purified *Mtb* σ^B^ *RNAP holoenzyme*

*Mtb* RNAP σ^B^ holoenzyme complex was purified from a 10X N-terminal His-tag on α??construct (pET-Duet-*rpoB*-*rpoC*, pAcYc-His*rpoA*-*rpoZ*, and pAC27-*sigB* plasmids) transformed in *E. coli* NiCo21(DE3) cells (NEB, C2529H, (116)). Cells were grown at 25°C to an OD_600_ of 0.6, induced with 0.5 mM IPTG and grown overnight at 16°C. Resuspension of pellets and protein purification was carried out identically to that described for *Mtb* RNAP core (Ni^2+^ HisTrap FF crude affinity followed by size exclusion chromatography), followed by anion exchange chromatography as described for the *Mtb* σ^A^ RNAP holoenzyme. The holoenzyme was dialyzed into storage buffer, concentrated and frozen as described for *Mtb* RNAP core, using an extinction coefficient of 264,955 M^-1^ cm^-1^.

#### Mtb σ^A^, Mtb σ^B^ and Mtb σ^A^_Δ205_

*Mtb* σ^A^ and *Mtb* σ^A^_Δ205_ were purified from pET-SUMO plasmid vectors transformed in *E. coli* BL21(DE3) and expressed as described for *Mtb* RNAP core above. Pellets were resuspended in 50 mM sodium phosphate, pH 8.0, 5 mM imidazole, 300 mM NaCl, 5 mM β-ME with 1 protease inhibitor cocktail tablet and lysed via sonication. After centrifugation (5,000 rpm for 30 min at 8°C), clarified lysate was incubated with Ni^2+^ NTA agarose beads (GoldBio, H350-50) for 30 min. The column was washed with 20 mM imidazole and protein was eluted with 250 mM imidazole in 50 mM sodium phosphate, pH 8.0, 300 mM NaCl. His-SUMO tag was cleaved by Ulp1 during overnight dialysis in 20 mM Tris, pH 8.0, 150 mM NaCl, 0.5 mM EDTA, 1 mM MgCl_2_, 5% (v/v) glycerol and 2 mM β-ME in 6–8 kDa MWCO tubing (Spectrum Labs, 132655). The σ factors and cleaved His-SUMO tag were then separated and further purified by anion exchange chromatography (MonoQ 5/50 GL, Cytiva, 17516601) using a 150–1000 mM NaCl gradient.

*Mtb* σ^B^ was purified similarly, but with the following changes. *E. coli* NiCo21(DE3) cells were used. Cell pellets were resuspended and sonicated in 50 mM potassium phosphate, pH 8.0, 20 mM imidazole, 500 mM ammonium chloride, 10% (v/v) glycerol, 0.1% triton X, 10 mM β-ME with 1 protease inhibitor cocktail tablet. Clarified lysate was loaded onto a Ni^2+^ HisTrap FF crude affinity column and protein was purified using a 5–1000 mM imidazole gradient in 20 mM potassium phosphate, pH 8.0 and 250 mM ammonium chloride. The dialysis buffer for Ulp1 cleavage was 20 mM potassium phosphate, pH 8.0, 250 mM ammonium chloride and 1 mM β-ME. The cleaved protein and the His-SUMO tag were then separated by an additional round of Ni^2+^ HisTrap FF crude affinity chromatography, where σ^B^ was collected from the flow-through.

All sigmas were then dialyzed into storage buffer as described for RNAP core and concentrated to ∼40 μM (5 kDa MWCO Vivaspin filters, Sartorius, F27335) using extinction coefficients of 35,410 M^-1^ cm^-1^ for *Mtb* σ^A^, 29,910 M^-1^ cm^-1^ for *Mtb* σ^A^_Δ205_, and 19,940 M^-1^ cm^-1^ for *Mtb* σ^B^.

#### Mtb CarD and RbpA

*Mtb* CarD and RbpA, in pET-SUMO plasmid vectors, were expressed, purified, and the His-SUMO tag was cleaved in accordance with methods previously described (27, 36). In some cases, after cleavage and separation of the SUMO tag, an additional anion exchange chromatography step (MonoQ 10/100 GL, Cytiva, 17516701) was implemented using a 100–1000 mM NaCl gradient. Eluted fractions were then dialyzed overnight in 20 mM Tris, pH 8.0, 150 mM NaCl, 1 β-ME, followed by concentration to ∼200 μM determined using extinction coefficients of 16,900 M^-1^ cm^-1^ for *Mtb* CarD and 13,980 M^-1^ cm^-1^ for *Mtb* RbpA.

### Plate-reader fluorescence measurements

For all experiments, data was collected using a CLARIOstar Plus Microplate reader (BMG LabTech) in a 384 well, low volume, round-bottom, non-binding polystyrene assay plate (Corning, 4514) with the corresponding Voyager analysis software. To measure multi-round and single-round transcription kinetics in real-time, we monitored the change in DFHBI fluorescence upon binding to a transcribed, full-length RNA sequence containing the iSpinach D5 aptamer. DFHBI fluorescence was measured with a monochromator excitation of 480 ± 15 nm, and the resulting emission signal was monitored at 530 ± 20 nm. To measure dissociation of promoter-bound complexes, Cy3 fluorescence was measured with a monochromator excitation of 535 ± 30 nm, and the resulting emission signal was monitored at 585 ± 30 nm. All reactions were at 10 μl final volume following initiation with 2.5 μl (single- and multi-round experiments) or 5 μl (dissociation experiments) from automated reagent injectors (BMG LabTech). Based on the volumes added for each corresponding buffer addition and concentrated stock component, the final solution conditions were 20 mM Tris (pH 8.0 at 37°C), 40 mM NaCl, 75 mM K-glutamate, 10 mM MgCl_2_, 5 μM ZnCl_2_, 20 μM EDTA, 5% (v/v) glycerol (defined as transcription buffer) with 1 mM DTT and 0.1 mg/ml BSA. All experiments were conducted at 37°C. For all experiments presented, independent preparations of *Mtb* holoenzymes were used.

#### Multi-round transcription experiments

Transcription reactions were performed with rNTPs (Thermo Scientific, R0481) at 1 mM, 3,5-difluoro-4-hydroxybenzylidene imidazolinone (DFHBI) dye (Sigma Aldrich, SML1627) at 20 μM, and RiboLock RNase inhibitors (Thermo Scientific, EO0381) at 0.4 U/μl. DFHBI concentration was determined using an extinction coefficient at 420 nm of 31,611 M^−1^ cm^−1^. When applicable, CarD and RbpA were added at concentrations of 1 μM and 2 μM, respectively. Transcription reaction master mixes contained 75% of the final volume which included the RNAP holoenzyme (co-expressed/purified or *in vitro* reconstituted), DFHBI, and RNase inhibitors, CarD/RbpA (when applicable), and either DNA or rNTPs, depending on how the reaction was initiated. For experiments comparing *rrnA* promoters and effects of DNA topology, master mixes containing 100 nM RNAP co-expressed/purified holoenzymes and 5 nM circular plasmid DNA templates along with all other components were preincubated for 15 min at 37°C before initiation with rNTPs. For σ titration experiments, 100 nM RNAP core was pre-incubated with various concentrations of σ for 15 min at 37°C along with all other reaction components before initiation with 5 nM *rrnA*P3 circular plasmid DNA template. Data was typically acquired in 10–20 s intervals, not exceeding 40 min total. A minimum of 3 technical replicates of the negative control (leaving out DNA or rNTP) were collected and measured concurrently with the experimental data. Using the average of this negative control, the experimental data was corrected as previously described (37), bringing all starting fluorescence values to zero and correcting for any time-dependent drift in fluorescence. Between 4 and 7 independent experiments were collected for each condition with 3 technical replicates each. Standard deviations were used as a statistical weight during the linear regression analyses as previously described to obtain the steady-state rate (37).

#### Single-round transcription experiments

To promote single-round conditions by preventing dissociated/terminated RNAPs from rebinding the promoter template, salmon-sperm DNA (Thermo Fisher Scientific, 15632011) was used as a competitor. Competitor DNA was buffer exchanged in transcription buffer and concentration was determined by A_260_. Promoter-bound complexes were pre-formed with 5 nM *rrnA*P3 circular plasmid DNA and either 250 nM σ^A^ RNAP co-expressed/purified holoenzyme supplemented with 2.5 μM σ^A^ or 125 nM σ^B^ RNAP co-expressed/purified holoenzyme supplemented with 1.25 μM σ^B^ along with 20 μM DFHBI and 0.4 U/μl RNase inhibitors. These concentrations were chosen to maintain the same protein:DNA ratio as used in the stopped-flow assays (below). Following a 15 min incubation at 37°C, reactions were initiated with 25 μg/ml of salmon-sperm DNA and 1 mM rNTPs. When competitor was first pre-incubated with RNAP holoenzyme and *rrnA*P3 circular plasmid DNA containing the aptamer sequence, no change in fluorescence upon initiating the reaction with NTPs was observed (**Figure 4B**). Data was acquired in 6 s intervals, for 20 min total and underwent the same subtractions as described for the multi-round experiments. The averages and standard deviations for 3 independent experiments are presented.

#### Dissociation of promoter-bound complexes

Dissociation of RNAP-promoter bound complexes experiments were adapted from previous methods described using stopped-flow rapid mixing (31) and those described below. Briefly, in a final volume of 10 µl, 100 nM σ^A^ RNAP co-expressed/purified holoenzyme or 50 nM σ^B^ RNAP co-expressed/purified holoenzyme supplemented with 500 nM σ^B^ and with 2 μM CarD/4 μM RbpA (when applicable) was incubated with 2 nM linear *rrnA*P3 promoter Cy3-labeled DNA at 37^?^C for 15 min. RNAP dissociation was measured by subsequent equal injection with 50 μg/ml salmon-sperm DNA. All concentrations listed are those prior to equal volume dilutions due to mixing. Data was acquired in 90 s intervals for a total of 20 hours. To avoid sample evaporation of the sample over time, immediately following injection of competitor DNA, wells were covered with an optical adhesive (Applied Biosystems, 4360954). Data was normalized to obtain the fraction of the initial signal remaining at a given time with the formula: [(F – F_o_)/(F_start_ – F_o_)] where F_o_ is the buffer subtracted value for Cy3-labeled DNA alone, F is the buffer subtracted signal, and F_start_ is the buffer subtracted signal at the time of injection. The averages and standard deviations for 3 independent experiments are presented, where the standard deviation was used as a statistical weight in fitting the time-courses to a single exponential function for the σ^A^ holoenzyme and a double exponential function for the σ^B^ holoenzyme.

### Stopped-flow fluorescence measurements

For all experiments, data was collected at 37°C using a SX-20 stopped-flow spectrophotometer (Applied Photophysics) with excitation provided by a 535 nm fixed-wavelength LED light source with a 550 nm short pass cut-off filter (Applied Photophysics), while monitoring emission using a 570 nm long pass cut-off filter (Newport Optics), as previously described (31). Following equal volume mixing with a total shot volume of 100 μl, the final reaction conditions corresponded to those of the transcription buffer with 1 mM DTT, and 0.1 mg/ml BSA. For all experiments presented, independent preparations of *Mtb* holoenzymes were used, where multiple technical replicates were combined to measure a ‘shot average’. The shot averages from each holoenzyme preparation were weighted equally in determining the averages and errors presented. Data was collected for 1000 or 2500 s with logarithmic sampling over 2500 points.

#### Open complex formation

For experiments comparing the basal kinetics of *Mtb* holoenzymes (**Figure 3A**), 50 nM σ^A^ RNAP co-expressed/purified holoenzyme and 50 nM σ^B^ RNAP co-expressed/purified holoenzyme supplemented with 500 nM σ^B^ were incubated at 37°C for 5 min prior to equal volume mixing with 2 nM linear *rrnA*P3 promoter Cy3-labeled DNA. Experiments in the presence of 2 μM CarD and 4 μM RbpA were conducted in the same way except that the σ^A^ RNAP co-expressed/purified holoenzyme concentration was increased to 100 nM to saturate the equilibrium fluorescence value (compare end point fluorescence value **Figure 3A** to **Figure 5A**). All concentrations listed are prior to equal volume mixing. Data is plotted as fold-change over DNA alone according to the formula: (F – F_o_)/F_o_, where F_o_ is the buffer subtracted value for Cy3-labeled DNA alone and F is the buffer subtracted signal.

#### Promoter escape

100 nM σ^A^ RNAP co-expressed/purified holoenzyme or 50 nM σ^B^ RNAP co-expressed/purified holoenzyme supplemented with 500 nM σ^B^ ± CarD/RbpA were pre-incubated with linear *Mtb rrnA*P3 promoter Cy3-labeled DNA and were rapidly mixed with 50 μg/ml salmon-sperm competitor DNA and 2 mM rNTPs (concentrations listed prior to equal volume mixing). As both the dissociation and promoter escape assays are conducted under single-round conditions, and as a result independent of RNAP holoenzyme concentration, these holoenzyme concentrations were chosen since the open-complex equilibration signal is saturated, both in the absence and presence of CarD/RbpA (**Figure 5A,B**).

#### Dissociation of promoter-bound complexes

Concentrations used for monitoring RNAP dissociation are identical to those described above for dissociation measurements using the plate-reader assay. Here an equal volume of 50 μg/ml salmon-sperm competitor DNA was mixed with the co-expressed/purified holoenzyme, and the linear *rrnA*P3 promoter Cy3-labeled DNA.

## Acknowldgements

The authors would like to thank Dr. Jerome Prusa for discussions at the outset of this work, Abby Tang for assistance with aptamer assay optimization, and Dr. Eric Tomko for critical reading of the manuscript.

## Authors contributions

Ana Ruiz Manzano: Conceptualization, Investigation, Methodology, and Writing—original draft. Drake Jensen: Conceptualization, Investigation, Methodology, Validation, and Writing. Eric Galburt: Conceptualization, Funding acquisition, Methodology, Supervision, Validation, Visualization and Writing—original draft.

