## Supplemental Information for "Regulation of Steady State Ribosomal Transcription in *Mycobacterium tuberculosis*: Intersection of Sigma Subunits, Superhelicity, and Transcription Factors"

**This PDF file includes:**

**Supplementary Figure 1**: Examples of fluorescence vs. time traces of both holoenzymes as a function of σ factor concentration under multi-round conditions.
**Supplementary Figure 2:** Effects of CarD and RbpA on the kinetics of open complex dissociation.
**Supplementary Figure 3:** Effects of CarD and RbpA on the kinetic half-lives of each σ factor-containing holoenzyme.
**Supplementary Figure 4:** Agarose gel showing the different DNA topologies obtained from the *rrnA*P3 plasmid DNA template.
**Supplementary Figure 5:** Gyrase treated DNA vs. miniprep steady-state rates.
**Supplementary Figure 6:** Dependence of steady-state transcription rates on topology and transcription factors for $\sigma^A$ and $\sigma^B$ holoenzymes.

**Supplementary Table 1.** $V_{max}$ and $K_m$ for σ factor titrations in the presence of CarD and RbpA.
**Supplementary Table 2.** Fold changes in steady-state transcription relative to no factors across differing DNA topologies.
**Supplementary Table 3.** Fold changes in steady-state transcription relative to supercoiled templates.
**Supplementary Table 4.** DNA non-template sequences used in real-time fluorescence transcription experiments.
**Supplementary Table 5.** Protein constructs used in the paper.

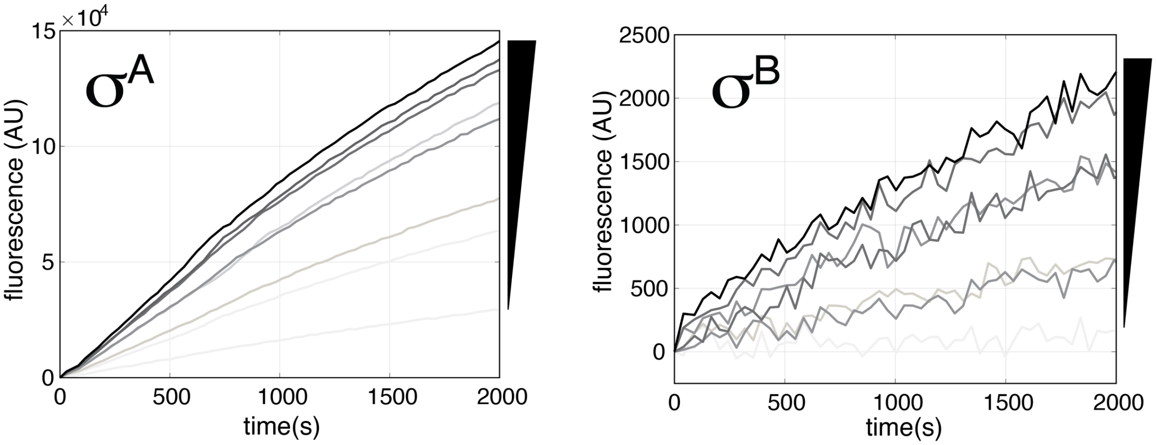


**Supplementary Figure 1**: **Examples of fluorescence vs. time traces of both holoenzymes as a function of σ factor concentration under *multi-round* conditions.** Increasing σ factor concentration titrated to 100 nM RNAP core is shown on the side and corresponds to darker traces. These raw data were averaged across four independent experiments and used for the quantification of steady-state transcription rates. Due to the initiation of the reaction via the addition of 5 nM *rrnA*P3 DNA, traces lack a burst phase (compare to **Figure 1C** in the main text which were initiated with 1 mM NTPs). Data were linearly fit starting from first measurement after injection of NTPs to extract rates.

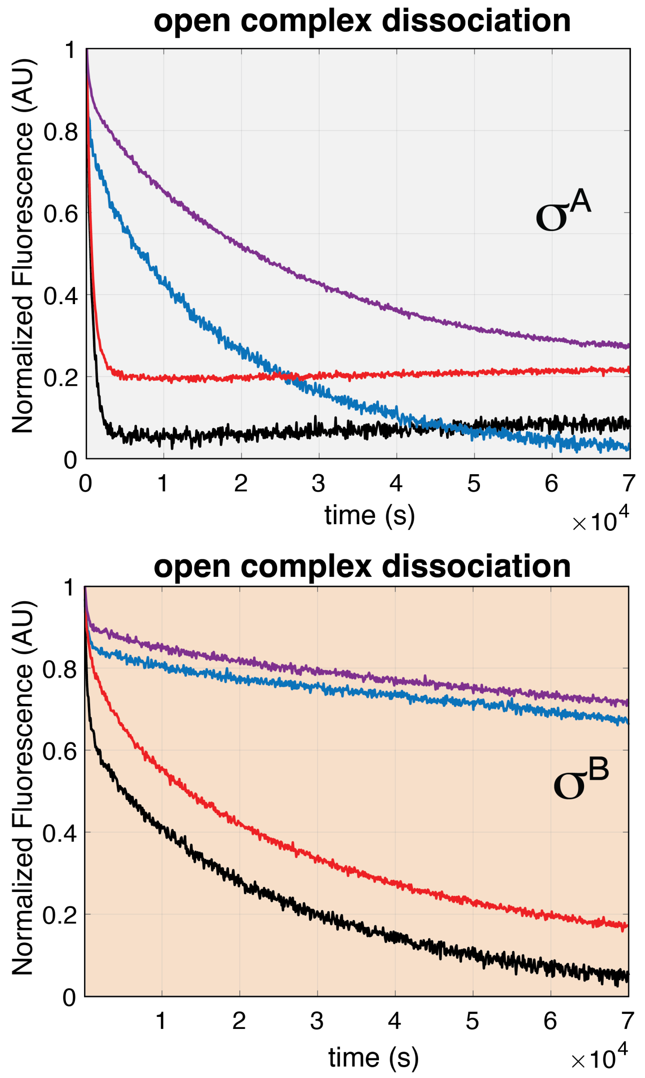


<u>**Supplementary Figure 2:**</u> **Effects of CarD and RbpA on the kinetics of open complex dissociation.** Plate reader dissociation data for both $\sigma^A$ (top panel, gray) and $\sigma^B$ (bottom panel, orange) holoenzymes collected in the absence of NTPs are shown with no factor (black), CarD (blue), RbpA (red), and both factors (purple). Holoenzyme complexes were pre-incubated with 2 nM linear *rrnA*P3 Cy3-labeled DNA and 50 µg/ml salmon-sperm DNA was introduced at time zero, leading to a time-dependent decrease in fluorescence signal. The averages from 3 independent experiments are plotted.

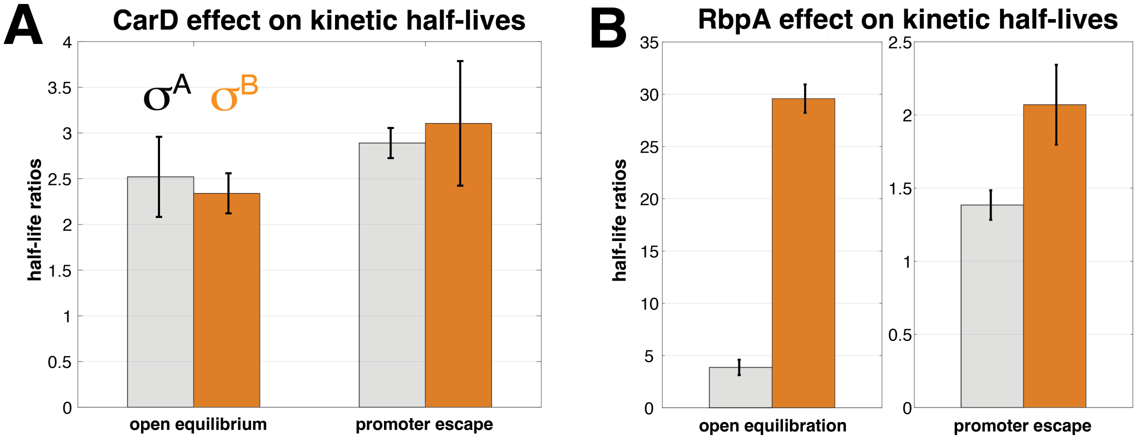


**<u>Supplementary Figure 3:</u> Effects of (A) CarD and (B) RbpA on the kinetic half-lives of each σ factor-containing holoenzyme.** The ratios of the half-lives for open complex equilibration and NTP-dependent decay in the context of $\sigma^A$ (grey) and $\sigma^B$ (orange). Half-lives were obtained from the time to reach half the total signal change from data presented in Figure 5. NTP-independent dissociation could not be compared as $\sigma^A$ holoenzyme lacks the longest time kinetic phase which the CarD and RbpA primarily effect. Each factor creates this phase de novo as can be seen in the traces themselves **(Supplementary Figure 2)**.

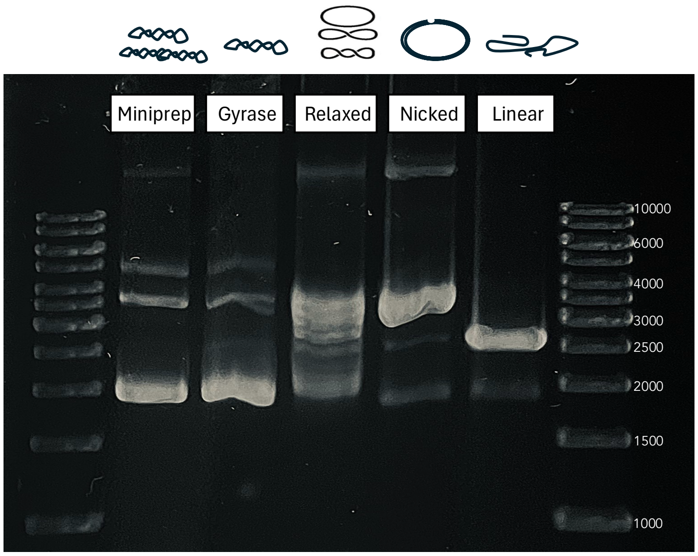


**<u>Supplementary Figure 4:</u> Agarose gel showing the different DNA topologies obtained from the *rrnA*P3 plasmid DNA template.** GeneRuler 1Kb DNA ladder (ThermoScientific, SM0313). Enzymes and conditions used to produce the different topological states can be found in the Methods.

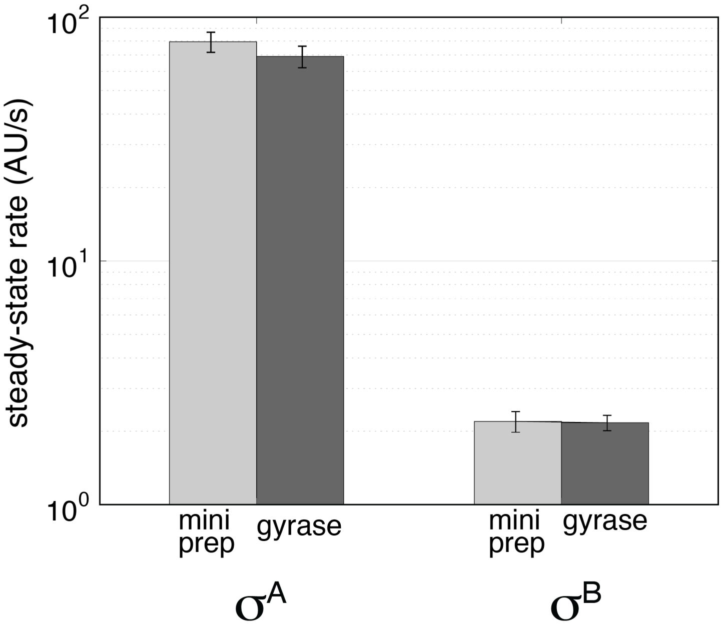


**<u>Supplementary Figure 5:</u> Gyrase treated DNA vs. miniprep steady-state rates.** Gyrase treated and miniprepped *rrnA*P3 plasmid DNA templates result in comparable steady-state rates suggesting a saturation of the topological effect for both $\sigma^A$ and $\sigma^B$ holoenzymes.

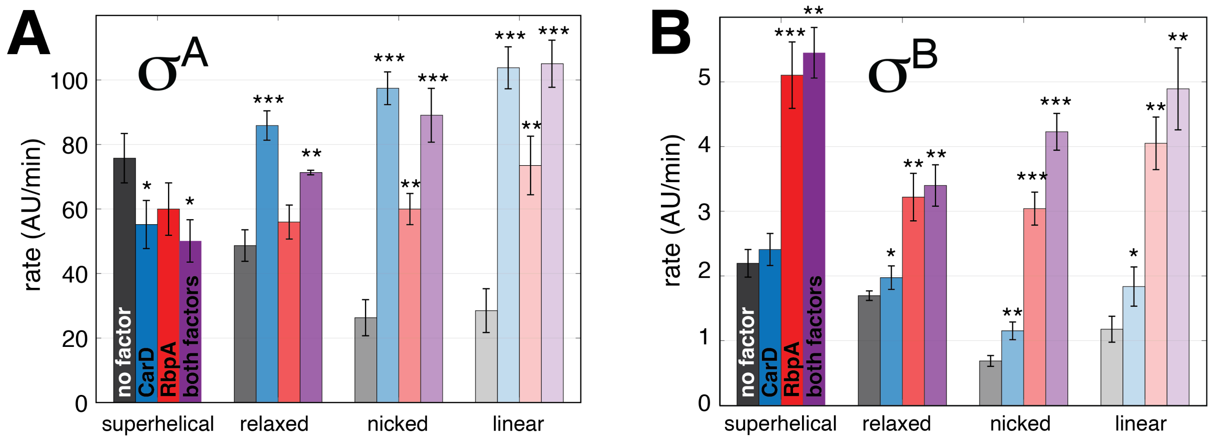


**<u>Supplementary Figure 6:</u> Dependence of steady-state transcription rates on topology and transcription factors for $\sigma^A$ (A) and $\sigma^B$ (B) holoenzymes.** Template topology is indicated on the x-axis and factor presence by bar color as follows: no factor (gray); CarD (blue); RbpA (red); and both factors (purple). Error bars indicate standard error of the mean from at least 6 traces in each condition collected across independent experiments, each with triplicate technical replicates. P-values (paired t-test) from the no factor condition are indicated by asterisks as follows: less than 0.05 (*), less than 0.005 (**), less than 0.0005 (***).

| Factor | Sigma | $K_m$ (nM) | | $V_{max}$ | |
|---|---|---|---|---|---|
| No factors | $\sigma^A$ | 193 | ± 50 | 93 | ± 7 |
| | $\sigma^B$ | 177 | ± 46 | 1.1 | ± 0.1 |
| | $\sigma^A$ Δ205 | 210 | ± 100 | 0.7 | ± 0.1 |
| CarD | $\sigma^A$ | 105 | ± 37 | 40 | ± 4 |
| | $\sigma^B$ | 230 | ± 125 | 1.3 | ± 0.3 |
| | $\sigma^A$ Δ205 | 146 | ± 90 | 1.1 | ± 0.2 |
| RbpA | $\sigma^A$ | 77 | ± 30 | 66 | ± 8 |
| | $\sigma^B$ | 260 | ± 100 | 1.7 | ± 0.3 |
| | $\sigma^A$ Δ205 | 118 | ± 80 | 1.2 | ± 0.3 |
| CarD+RbpA | $\sigma^A$ | 56 | ± 17 | 41 | ±2 |
| | $\sigma^B$ | 254 | ± 95 | 1.5 | ± 0.4 |
| | $\sigma^A$ Δ205 | 119 | ± 80 | 1.5 | ± 0.3 |

**Supplementary Table 1: $K_m$ and $V_{max}$ from σ factor titration fits in the presence of CarD and RbpA**. Data calculated from the steady-state transcription rates as a function of σ factor concentration fit to hyperbolic curves. Errors indicate 95% confidence from fits of averaged data. See Figures 2 and 7 for corresponding data.

| | | $\sigma^A$-RNAP | | | | | $\sigma^B$-RNAP | | | | |
|---|---|---|---|---|---|---|---|---|---|---|---|
| | | **Supercoiled** | **Gyrase** | **Relaxed** | **Nicked** | **Linear** | **Supercoiled** | **Gyrase** | **Relaxed** | **Nicked** | **Linear** |
| CarD | **CarD/no factor** | 0.73 | 0.74 | 1.76 | 3.71 | 3.64 | 1.10 | 1.06 | 1.16 | 1.68 | 1.77 |
| | *P VALUE* | *0.01* | *0.06* | *8E-04* | *2.6E-06* | *2.0E-04* | *0.11* | *0.06* | *0.05* | *0.003* | *0.03* |
| RbpA | **RbpA/no factor** | 0.79 | 0.67 | 1.15 | 2.28 | 2.58 | 2.32 | 1.98 | 1.90 | 4.43 | 3.89 |
| | *P VALUE* | *0.2713* | *0.0058* | *0.2787* | *0.0012* | *0.0006* | *0.0005* | *0.0003* | *0.0012* | *0.0001* | *0.0008* |
| CarD + RbpA | **factors/no factor** | 0.66 | 0.71 | 1.47 | 3.39 | 3.69 | 2.48 | 1.96 | 2.00 | 6.16 | 4.70 |
| | *P VALUE* | *0.03* | *0.06* | *0.005* | *0.00014* | *1.97E-05* | *0.00115* | *0.005* | *0.002* | *2E-05* | *0.0021* |

**Supplementary Table 2: Fold changes in steady-state transcription relative to no factors across differing DNA topologies.** P-values were calculated using the paired t-test. See Figure 6 and Supplemental Figure 6 for corresponding data.

| **Signal over supercoiled** | $\sigma^A$-RNAP | | | | $\sigma^B$-RNAP | | | |
|---|---|---|---|---|---|---|---|---|
| | **no factor** | **CarD** | **RbpA** | **CarD + RbpA** | **no factor** | **CarD** | **RbpA** | **CarD + RbpA** |
| Gyrase | 0.91 | 0.92 | 0.77 | 0.97 | 0.99 | 0.96 | 0.84 | 0.78 |
| *P VALUE* | *0.05* | *0.89* | *0.14* | *0.518* | *0.82* | *0.71* | *0.15* | *0.135* |
| Relaxed | 0.64 | 1.56 | 0.93 | 1.42 | 0.77 | 0.82 | 0.63 | 0.62 |
| *P VALUE* | ***0.021*** | ***0.01*** | *0.63* | *0.052* | *0.1* | *0.05* | *0.02* | *0.02* |
| Nicked | 0.35 | 1.77 | 1.0 | 1.78 | 0.31 | 0.48 | 0.6 | 0.78 |
| *P VALUE* | *0.0016* | *0.002* | *1.0* | *0.017* | *0.007* | *0.01* | *0.004* | *0.059* |
| Linear | 0.38 | 1.88 | 1.23 | 2.1 | 0.47 | 0.76 | 0.79 | 0.9 |
| *P VALUE* | *0.0006* | *0.002* | *0.09* | *0.001* | *0.04* | ***0.01*** | *0.08* | *0.356* |

**Supplementary Table 3: Fold changes in steady-state transcription relative to supercoiled templates.** P-values were calculated using the paired t-test. See Figure 6 and Supplemental Figure 6 for corresponding data.

| Template Name | Insert Length | Construct Length | Insert Sequence |
|---|---|---|---|
| **AP1_2T plasmid** | 380 | 2601 | CGATCTAGAGTCTACCGGCCACCAGACATGATGGGCGCACCAGCGCCCATCATGTTCTTTTCGAAAGCTTCGGCCCG<br>ACCCCACATCGATCGCGGATATCCGTTGTTCGTGGAGAACCTGGTGAGTCTCGGTGCCGAGATCGAACGGGTATGC<br>TGTTAGGC**G**ACGGTCACCTATGGGCGGAAACAAGCAAGC**GGGAGTACGGTGAGGGTCGGGTCCAGTAGCTTCGG**<br>**CTACTGTTGAGTAGAGTGTGGGCTCCGTACTCCC**GTGTTGTTTGAGAACTCAATAGTGTGTTTGGTGGTTTCACCAG<br>GCATCAAATAAAACGAAAGGCTCAGTCGAAAGACTGGGCCTTTCGTTTTATCTGTTGTTTGTCGGTGAACGCTCTC |
| **AP1AP3_2T plasmid** | 408 | 2629 | CGATCTAGAGTCTACCGGCCACCAGACATGATGGGCGCACCAGCGCCCATCATGTTCTTTTCGAAAGCTTCGGCCCG<br>ACCGGTGCCGAGATCGAACGGGTATGCTGTTAGGCGACGGTCACCTATGGATATCTATGGATGACCGAACCTGGTC<br>TTGACTCCATTGCCGGATTTGTATTAGACTGGCAGG**G**TTGCCCCGAAGCGGGCGGAAACAAGCAAGC**GGGAGTAC**<br>**GGTGAGGGTCGGGTCCAGTAGCTTCGGCTACTGTTGAGTAGAGTGTGGGCTCCGTACTCCC**GTGTTGTTTGAGAA<br>CTCAATAGTGTGTTTGGTGGTTTCACCAGGCATCAAATAAAACGAAAGGCTCAGTCGAAAGACTGGGCCTTTCGTTT<br>TATCTGTTGTTTGTCGGTGAACGCTCTC |
| **AP3_2T plasmid** | 380 | 2601 | CGATCTAGAGTCTACCGGCCACCAGACATGATGGGCGCACCAGCGCCCATCATGTTCTTTTCGAAAGCTTCGGCCCG<br>ACCAGGCGACGGTCACCTATGGATATCTATGGATGACCGAACCTGGTCTTGACTCCATTGCCGGATTTGTATTAGAC<br>TGGCAGG**G**TTGCCCCGAAGCGGGCGGAAACAAGCAAGC**GGGAGTACGGTGAGGGTCGGGTCCAGTAGCTTCGG**<br>**CTACTGTTGAGTAGAGTGTGGGCTCCGTACTCCC**GTGTTGTTTGAGAACTCAATAGTGTGTTTGGTGGTTTCACCAG<br>GCATCAAATAAAACGAAAGGCTCAGTCGAAAGACTGGGCCTTTCGTTTTATCTGTTGTTTGTCGGTGAACGCTCTC |
| **PL _2T plasmid** | 380 | 2601 | CGATCTAGAGTCTACCGGCCACCAGACATGATGGGCGCACCAGCGCCCATCATGTTCTTTTCGAAAGCTTCGGCCCG<br>ACCATGTATAGTCTGCGCACAGTACTGCTTTCATAGTGAAGAACAGCTCTGCCCAGTACCAGACGTAGGTGGTTCCA<br>TCGGGTTACGGCCTCGGGGTGGGCGGAAACAAGCAAGCG**GGGAGTACGGTGAGGGTCGGGTCCAGTAGCTTCG**<br>**GCTACTGTTGAGTAGAGTGTGGGCTCCGTACTCCC**GTGTTGTTTGAGAACTCAATAGTGTGTTTGGTGGTTTCACCA<br>GGCATCAAATAAAACGAAAGGCTCAGTCGAAAGACTGGGCCTTTCGTTTTATCTGTTGTTTGTCGGTGAACGCTCTC |
| **AP3 Linear** | N/A | 150 | GGCGACGGTCACCTATGGATATCTATGGATGACCGAACCTGGTCTTGACTCCATTGCCGGATTTGTATTAGACTGGC<br>AGG**G**TTGCCCCGAAGCGGGCGGAAACAAGCAAGCGTGTTGTTTGAGAACTCAATAGTGTGTTTGGTGGTTTCA |

**Supplementary Table 4: DNA non-template sequences used in real-time fluorescence transcription experiments.** All templates used for single- and multi-round transcription experiments contain: a tuf Terminator (brown), promoter sequence spanning from -81 to +15 relative to the transcription start site, +1 in bold. Genomic sequence found downstream of the *rrnA*P3 promoter from+15 to +31, spinach i-spD5 (green), genomic sequence found downstream of the *rrnA*P3 promoter from +32 to +70, and the *rrnB* T1 Terminator (brown). 2T denotes the presence of both upstream and downstream terminators relative to the promoter. PL denotes Promoter Less plasmid. The AP3 linear template used for open complex equilibration, dissociation, and promoter escape experiments contains the *Mtb rrnA*P3 genomic DNA from -80 to + 70. Either specify or change color of the purple sequences before the aptamer in P1/P3, P3, and PL.

| Recombinant proteins | plasmids | Reference |
|---|---|---|
| ***Mtb* RNAP core** | | |
| pβ-β', pHis$\alpha$-$\omega$ | pETDuet-**rpoC/rpoB** | J. Mukhopadhyay |
| | pAcYc-**His_rpoA/rpoZ** | J. Mukhopadhyay |
| ***Mtb* RNAP $\sigma^A$** | | |
| pβ-β', pHis$\alpha$-$\omega$, p$\sigma^A$ | pETDuet-**rpoC/rpoB** | J. Mukhopadhyay |
| | pAcYc-**His_rpoA/rpoZ** | J. Mukhopadhyay |
| | pAC27-**sigA** | This paper |
| pβ-β', p$\alpha$-His$\sigma^A$, p$\omega$ | pETDuet-**rpoC/rpoB** | J. Mukhopadhyay |
| | pAcYcDuet-**His_sigA/rpoA** | J. Mukhopadhyay |
| | pCDF-**rpoZ** | J. Mukhopadhyay |
| ***Mtb* RNAP $\sigma^B$** | | |
| pβ-β', pHis$\alpha$-$\omega$, p$\sigma^B$ | pETDuet-**rpoC/rpoB** | J. Mukhopadhyay |
| | pAcYc-**His_rpoA/rpoZ** | J. Mukhopadhyay |
| | pAC27-**sigB** | This paper |
| ***Mtb* $\sigma^A$** | | |
| | pET-SUMO-**sigA** | This paper |
| ***Mtb* $\sigma^B$** | | |
| | pET-SUMO-**sigB** | This paper |
| ***Mtb* $\sigma^A\Delta 205$** | | |
| | pET-SUMO-**sigA_Δ205** | This paper |
| ***Mtb* CarD** | | |
| | pET-SUMO-**carD** | C. Stallings |
| ***Mtb* RbpA** | | |
| | pET-SUMO-**rbpA** | C. Stallings |

**Supplementary Table 5: Protein constructs used in the paper.** RNAP holoenzymes were assembled in two different ways: 1) co-expression and purification with all subunits from the same bacterial strain, and 2) a reconstituted version purifying RNAP core and SUMO-cleaved $\sigma$ subunits separately.
